# Lyso-Phosphatidic Acid Acyl-Transferases: a link with intracellular protein transport in *Arabidopsis* root cells?

**DOI:** 10.1101/2021.06.08.447563

**Authors:** Valérie Wattelet-Boyer, Marina Le Guédard, Franziska Dittrich-Domergue, Lilly Maneta-Peyret, Verena Kriechbaumer, Yohann Boutté, Jean-Jacques Bessoule, Patrick Moreau

## Abstract

Phosphatidic acid (PA) and Lysophosphatidic acid acyltransferases (LPAATs) might be critical for the secretory pathway. Four extra-plastidial LPAATs (numbered 2,3,4 and 5) were identified in *A. thaliana*. These AtLPAATs, displaying an enzymatic activity specific for LPA to produce PA, are located in the endomembrane system. We focused on the putative role of the AtLPAATs 3, 4 and 5 in the secretory pathway of root cells through genetical (knock-out mutants), biochemical (activity inhibitor, lipid analyses) and imaging (live and immuno-confocal microscopies) approaches. Treating a *lpaat4;lpaat5* double mutant with the LPAAT inhibitor CI976 showed a primary root growth decrease. The transport of the auxin transporter PIN2 was disturbed in this *lpaat4;lpaat5* double mutant treated with CI976, but not that of H+-ATPases. The *lpaat4;lpaat5* double mutant was sensitive to salt stress and the transport of the aquaporin PIP2;7 to the plasma membrane in the *lpaat4;lpaat5* double mutant treated with CI976 was reduced. We measured the amounts of neo-synthesized PA in roots, and found a decrease in PA only in the *lpaat4;lpaat5* double mutant treated with CI976, suggesting that the protein transport impairment was due to a critical PA concentration threshold.

**Highlight:** Phosphatidic acid produced by Lyso-Phosphatidic Acid Acyl-Transferases has an impact on the efficiency of the intracellular transport of some proteins in *Arabidopsis thaliana* root cells.

## Introduction

Lipids3 are very critical for organelle compartmentalization and membrane domain partition in all eukaryotic cells. In plant cells we, and other teams, have evidenced the involvement of lipids and their metabolism in the regulation of the plant secretory pathway (Melser et al., 2011; Boutté and Moreau, 2014). All lipid families appear to be involved as sterols (Laloi et al., 2007; Men et al., 2008; Boutté et al., 2010), sphingolipids (Melser et al., 2010; Markham et al., 2011; Wattelet-Boyer et al., 2016) and glycerolipids (Pleskot et al., 2012; Boutté and Moreau, 2014). Lipids play critical roles in endomembrane morphodynamics regulation, organelle morphology and transport vesicle formation/fusion as indicated with *in vivo/in vitro* studies in various eukaryotic models (Yang et al., 2008, 2011; Ha et al., 2012; Boutté and Moreau, 2014; Melero et al., 2018).

A role for several enzymes of phospholipid metabolism such as phospholipases and lysophospholipid acyltransferases (LPATs) for the function of the secretory/retrograde pathways has been particularly highlighted in animal and yeast cells (Yang et al., 2008, 2011; Melero et al., 2018). In plant cells, several phospholipases have been shown to be required for the functionality of the secretory pathway (Li et al., 2007; Lee et al., 2010; Li et al., 2011; Kim et al., 2011). However, behind their expected role in phospholipid biosynthesis, no link has been yet revealed between LPATs and the functionality of the secretory pathway in plant cells. Among lysophospholipid acyltransferases, lysophosphatidic acid (LPA) acyltransferases (LPAATs) may be very critical as phosphatidic acid (PA) and LPA have been shown to have an important impact on the functionality of the secretory pathway in animal cells (Yang et al., 2008, 2011). Therefore, PA and its precursor LPA, in addition to their role as precursors of *de novo* phospholipid biosynthesis and their known involvement in many signalling pathways (Pokotylo et al., 2018; Yao and Xue, 2018), deserve our interest in possible functions in the secretory pathway in plant cells.

The amount of PA in the cell depends on the *de novo* synthesis via the Kennedy pathway but can also result from the activity of multiple enzymes such as phospholipases D, diacylglycerol kinases or LPAATs that will increase the PA pool. In contrast, PA phosphatases or phospholipases A1/A2 cause a decrease in the amount of PA in the cells. A study carried out in *Nicotiana tabacum* indicated that pharmacological inhibition of most of these enzymes leads to very different effects on pollen tube growth (Pleskot et al., 2012), indicating that there could be different PA pools related to different enzyme activities. At present, no studies have been conducted specifically to investigate the putative role of LPAATs in the secretory pathway of plant cells. We hypothesize that LPAATs could, as shown for phospholipases, participate in the regulation of membrane curvature by catalyzing PA production from LPA (two molecules with different physicochemical properties) and therefore contribute to the regulation of membrane trafficking (Yang et al., 2008, 2011; Boutté and Moreau, 2014).

Several membrane-bound LPAATs have been identified so far in *A. thaliana* (Kim and Huang, 2004; Kim et al., 2005; Wang et al., 2013; Körbes et al., 2016; Angkawijaya et al., 2017; Angkawijaya et al., 2019). In higher eukaryotes, these enzymes were either named LPAT or LPAAT but we prefer and propose to use the name LPAAT to precisely highlight their lysophosphatidic acid acyltransferase activities. It is effectively inconsistent that the first identified isoform was named “LPAAT1” whereas the following isoforms were reported as “LPAT2-5” (Kim et al 2005). AtLPAAT1 has been found to be involved in *de novo* synthesis of PA in the plastids (Kim and Huang, 2004; Yu et al., 2004). The ER-located AtLPAAT2 (LPAT2) has been shown to be critical for female gametophyte development in Arabidopsis (Kim et al., 2005). In addition, the over-expression of AtLPAAT2, which stimulates the *de novo* production of phospholipids, resulted in the enhanced primary root growth in phosphate-starved *Arabidopsis* (Angkawijaya et al., 2017). This likely indicates AtLPAAT2 as a primordial enzyme for the *de novo* synthesis of PA in the ER. Three other potential LPAAT genes, called *AtLPAAT3, AtLPAAT4* and *AtLPAAT5*, have been identified. Recently, Angkawijaya et al. (2019) have shown that AtLPAAT4 and AtLPAAT5 can be involved in the neo-synthesis of phospholipids and triglycerides in response to nitrogen starvation. Since AtLPAAT2 is probably the main source of PA for the *de novo* synthesis of phospholipids (Kim et al., 2005; Angkawijaya et al., 2017), we have investigated whether the other AtLPAATs can be associated with a further role in membrane dynamics linked to the functioning of the secretory pathway.

We first showed that these AtLPAATs have an enzymatic activity specific for producing PA from lysophosphatidic acid and that they are located in the endomembrane system (mainly ER). Through genetic, biochemical and imaging approaches, we show that a *lpaat4;lpaat5* double mutant, sensitive to salt stress, presents a primary root growth decrease phenotype when treated with the LPAT inhibitor CI976 (Brown et al., 2008; Schmidt and Brown, 2009; Yang et al., 2011) and that the transport of the aquaporin PIP2;7 is affected. In addition, the transport of the auxin carrier PIN2 was also disturbed. By measuring the amounts of PA in the roots, we were able to link the disturbance of protein transport to a critical PA concentration threshold.

## Material and Methods

### *Arabidopsis* material and growth conditions

The *Arabidopsis thaliana* ecotype Colombia-0 (Col-0) and the following mutants were used: *lpaat3-1* (SALK_046680), *lpaat4-2* (GK_899A04) and *lpaat5-2* (SALK_020291). Double mutants *lpaat3-1;lpaat4-2, lpaat3-1;lpaat5-2* and *lpaat4-2;lpaat5-2* were obtained crossing the previously listed lines. Triple mutant *lpaat3-1;lpaat4-2;lpaat5-2* was obtained crossing the double mutant *lpaat4-2;lpaat5-2* and SALK_046680. The following pPIN2::PIN2–GFP (Xu et al., 2005) transgenic fluorescent protein marker line in Col-0 was used for crossing with double mutant *lpaat4-2;lpaat5-2*.

Seeds were sterilized by treatment with 95% (v/v) ethanol for 10 seconds, followed by bleach solution for 20 minutes, then repeatedly washed with sterile water. Seeds were then sown on half Murashige and Skoog (MS) agar medium plates (0.8% plant agar (Meridis #P1001,1000), 1% sucrose (Merck # 84100) and 2.5□mM morpholinoethanesulfonic acid (Euromedex # EU0033) pH 5.8 with KOH, left at 4□°C for 2 days and then grown vertically in 16□h light/8□h darkness for 5 days before all experiments.

### Inhibitor treatment

CI976 was used as a lysophosphatidyl acid acyltransferase inhibitor (Merck # C3743). CI976 intermediate stock solution was made at 10mM in dimethylsulfoxide (DMSO), stocked at - 20°C and added in media prepared extemporarily. Seedlings were grown on half MS plate containing 10µM of the inhibitor in most of the experiments, except to determine the WT, double mutant *lpaat4-2;lpaat5-2* and triple mutant *lpaat3-1;lpaat4-2;lpaat5-2* lines sensitivity to the treatment, where 5µM of inhibitor was also tested. In all experimental conditions, the final DMSO concentration was the same for the controls without CI976 and at the different concentration of CI976 used.

### Phenotypical characterization

Seedlings were grown on half MS agar medium plate containing various CI976 concentrations (0, 5 or 10 µM). Root length was measured five days after germination using the ImageJ software. To compare experiments, all the measured values were standardized to the WT median value in untreated condition for each experiment.

To assess the different lines sensitivity to salt stress, 50 mM NaCl (Euromedex # 1112) was added to the half MS agar medium.

### Cloning and transgenic plants

Sequence data of AtLPAATs can be found in the Arabidopsis Genome Initiative or GenBank/EMBL databases under the following accession numbers: *AtLPAAT2*, At3g57650; *AtLPAAT3*, At1g51260; *AtLPAAT4*, At1g75020; *AtLPAAT5*, At3g18850. Coding sequence of *AtLPAAT2, AtLPAAT4* were amplified on leaves cDNA, while *AtLPAAT3* and *AtLPAAT5* were amplified on flowers cDNA. We used respectively the primer pairs P1531/P1533, P1539/P1541, P1535/P1537 and P1543/P1545. To generate the following diK mutants: *AtLPAAT2 K387A/K389A, AtLPAAT4 K374A/K376A* and *AtLPAAT5 K371A/K375A* we used respectively the following primer pairs containing the mutation: P5294/P5431, P2051/P2052 and P5292/P5432. *LPAAT3* diacidic mutant 1 (D74G/A75/E76G) and 2 (D293G/L294A/E295G) were obtained by overlapping PCR using respectively primers P2578/P2580/P2581/P2582/P66/P67 and P2578/P2580/P2585/P2586/P66/P67. Amplified sequences were cloned by BR reactions in entry vectors pDONR™221 or pENTR-d-TOPO™ (Thermofisher Scientific) using Gateway^®^ recombinational cloning technology (Thermofisher Scientific). For expression in *E*.*coli*, entry vectors were cut by NcoI (Biolabs #R0193) and XhoI (Biolabs #R0146) restriction endonucleases. The product was cloned into the pET-15b vector (Novagen). For expression in plant, AtLPAAT entry vectors and pK7WGF2 destination vectors were combined by LR recombination using Gateway^®^ recombinational cloning technology (Thermofisher Scientific).

To complement the double mutant *lpaat4-2;lpaat5-2* we generated the construct p*AtLPAAT4*:tagRFP-*AtLPAAT4g*. For that, we amplified the *AtLPAAT4* promoter sequence and the *AtLPAAT4* full length DNA genomic sequence using respectively the primer pairs P5730/P5731 and P5736/5737. Each PCR product was purified and cloned respectively in pDONR™ P4-P1r vector and pDONR™ P2r-P3 vector (Thermofisher Scientific). We used a third vector containing tagRFP in pDONR™221 backbone (Thermofisher Scientific) and generated the final construct in the pH7m34GW destination vector using the Multisite Gateway® cloning system (Thermofisher Scientific).

All PCR amplifications were performed using Q5™ High-Fidelity DNA polymerase at the annealing temperature and extension times recommended by the manufacturer (Biolabs #M04915). PCR fragments and plasmids were respectively purified with NucleoSpin^®^ Gel and PCR cleanup (Macherey-nagel # 740609) and NucleoSpin^®^ Plasmid (Macherey-Nagel #740499). All the entry vectors were sequenced, and sequences were analyzed with CLC Mainwork Bench 6. Primers used in this study are listed in Supplementary Table S1.

For transient expression in *Arabidopsis* cotyledons, seeds were sterilized as described above and sown in 6 wells culture plates containing 4 ml half MS agar medium. Plates were left at 4□°C for 2 days and then grown in 16□h light/8□h darkness for 7 days.

Constructs were transferred into the *Agrobacterium tumefaciens* C58C1Rif^R^ strain harboring the plasmid pMP90. 4 days after germination *Agrobacterium tumefaciens* suspension in MS-Glu liquid medium (0.21% MS (w/v), 2% glucose (w/v), 0.39% MES (w/v), 0.05%Tween, 200mM acetosyringone, pH5.7) was used to transform transiently the cotyledons. For that, seedlings were incubated 40 min at RT with suspension of *Agrobacterium tumefaciens* expressing natives or mutant LPAATs and HDEL marker at 1 OD_600nm_ and 0.2 OD_600nm_ respectively. Suspension was then removed and plate were left in 16□h light/8□h darkness until the 7^th^ day after germination.

To study the effect of the double mutation *lpaat4-2;lpaat5-2* on PIN2-GFP subcellular localization at the plasma membrane, the double mutant *lpaat4-2;lpaat5-2* and the pPIN2::PIN2–GFP.transgenic line (Xu *et al*., 2005) were crossed. Primer pairs 1905/1906 (LP/RP SALK-020291), 5744/5745 (LP/RP GABI_899A04) and LBa1 were used for genotyping on ammonium glufosinate (10µg/ml) resistant seedlings.

### RNA extraction, RT-PCR and qPCR

Tissues were disrupted using stainless steel beads 5mm (Qiagen#69989) and Tissuelyser II (Qiagen). Total RNA were extracted from roots 5 days after germination using the RNeasy^®^ Plant Mini kit (Qiagen #74904) according to the manufacturer’s instructions. First strand cDNA was synthesized using SuperScript^®^ II Reverse Transcriptase (ThermoFisher # 18064014) and OligodT. Then, mRNAs were treated with DNase I using DNa-*free*™ Kit (ThermoFisher # AM1906). Expression analysis of *AtLPAAT2, AtLPAAT3, AtLPAAT4* and *AtLPAAT5* by RT-qPCRs were performed with the Bio-Rad CFX96 real-time system using GoTaq^®^ qPCR Master mix (Promega # A6002). The specific primer pairs used for *AtLPAAT2, AtLPAAT3, AtLPAAT4, AtLPAAT5, EF-1a* and *At4g33380* were P5412/P5413, P5414/P5415, P5783/P5784, P5418/P5419, P4833/P4834 and P4847/P4848, respectively. The transcript abundance in samples was determined using a comparative cycle threshold (*C*_t_) method. The relative abundance of *EF-1a and At4g33380* mRNAs in each sample was determined and used to normalize for differences of total RNA level. Semi-quantitative RT-PCR analysis of steady-state At*LPAAT*s transcripts in roots from 5 days old plants was performed to compare *lpaat4-2;lpaat5-2* and *lpaat3-1;lpaat4-2;lpaat5-2* mutant lines with the wild-type plants. The *EF-1a* gene was used as a constitutively expressed control. We used GoTaq® G2 DNA Polymerase (Promega). All primers are listed in Supplementary Table S1, the characterisation of *lpaat* insertion mutant lines and T-DNA positions is shown in Supplementary Fig. S1, and controls for *lpaat4-2;lpaat5-2* and *lpaat3-1;lpaat4-2;lpaat5-2* mutant lines are shown in Supplementary Fig. S2.

### Immunocytochemistry, FM4-64 uptake and confocal laser scanning microscopy

Whole-mount immunolabelling of *Arabidopsis* root was performed as described (Boutté and Grebe, 2014). In brief, 5 day-old seedlings were fixed in 4% paraformaldehyde dissolved in MTSB (50□mM PIPES, 5□mM EGTA, 5□mM MgSO_4_ pH 7 with KOH) for one hour at room temperature (RT) and washed three times with MTSB. Roots were cut on superfrost slides (Menzel Gläser, Germany) and dried at RT. Roots were then permeabilized with 2% Driselase (Merck #D9515), dissolved in MTSB for 30□min at RT, rinsed four times with MTSB and treated for one hour at RT with 10% dimethylsulfoxide + 3% Igepal CA-630 (Merck # I3021) dissolved in MTSB. Aspecific sites were blocked with 5% normal donkey serum (NDS, Merck # D9663) in MTSB for one hour at RT. Primary antibodies, in 5% NDS/MTSB, were incubated overnight at 4□ °C and then washed four times with MTSB. Secondary antibodies, in 5% NDS/MTSB, were incubated one hour at RT and then washed four times with MTSB.

Primary antibodies were diluted as follows: rabbit anti-PIP2;7 1:400 (Agrisera, AS09469), rabbit anti-H+ATPase 1:1000 (Agrisera AS07260), rabbit anti-echidna 1:600 (Gendre et al., 2011 and Boutté et al., 2013), rabbit anti-Membrine11 1:300 (Marais et al., 2015), rabbit anti-SAR1 1:250 (Agrisera AS08326). Dilution of secondary antibody donkey anti-rabbit IgG AlexaFluor 488-coupled (Jackson Immunoresearch, 711-605-152) was 1:300.

FM4-64 uptake was performed on 5 days old seedling that had grown on half MS agar media containing 10 µM of CI976. Seedlings were dark incubated in 5 µM FM4-64 solution for 5 min. Rapid wash in half MS solution containing 10 µM of CI976 followed, and FM4-64 uptake was determined each minutes on root epidermal cells by confocal microscopy from 3 to 15 min after seedlings wash.

Confocal laser scanning microscopy was performed using Zeiss LSM880. For live-cell imaging, seedlings were mounted with MS liquid medium. Laser excitation lines for the different fluorophores were 488 □ nm for GFP, AlexaFluor 488 and FM4-64, and 561 □ nm for mCherry. Fluorescence emissions were detected at 490–570 □ nm for GFP and AlexaFluor 488, 620-695 nm for FM4-64 and 580–660□nm for mCherry. Scanning was performed with a pixel dwell of 3µs. In multi-labelling acquisitions, detection was in sequential line-scanning mode. An oil-corrected × 63 objective, numerical aperture=1.4 (C Plan Apochromat 63.0×1.40 OIL DIC UV VIS IR-M27) was used in immunolabelling and live-cell imaging experiments, except for FM4-64 uptake where an oil-corrected × 40 objective, numerical aperture=1,3 (Plan Apochromat 40.0×1.3 OIL DIC UV IR-M27) was used. Image analysis was performed using ZEN lite 2.6 2018 (Zeiss) and ImageJ softwares.

### Tobacco leaf infiltration and confocal microscopy

*Nicotiana tabacum* (SR1 cv Petit Havana) plants were grown in a greenhouse for transient expression of fluorescent constructs according to Sparkes et al. (2006). In brief, each expression vector was introduced into the *Agrobacterium tumefaciens* strain GV3101 by heat shock transformation. Transformed colonies were inoculated into 5 ml of YEB medium (5 g/l beef extract, 1 g/l yeast extract, 5 g/l sucrose and 0.5 g/l MgSO_4_·7H_2_O) with 50 μg/ml spectinomycin and rifampicin. The bacterial culture was incubated overnight in a shaker at 180 rpm 25 °C. 1 ml of the bacterial cultures was pelleted by centrifugation at 1800 g at room temperature for 5 minutes. Pellets were washed with infiltration buffer (5mg/ml glucose, 50mM MES, 2mM Na_3_PO_4_·12H_2_O and 0.1 mM acetosyringone) and then resuspended also in infiltration buffer. The bacterial suspension was diluted in infiltration buffer to a final OD_600_ of 0.1 for each construct. The bacterial solution was injected into the underside of the tobacco leaf using a 1ml syringe. Infiltrated plants were incubated at 22°C for three days prior to imaging. Leaf epidermal samples were imaged using a Zeiss PlanApo 100×/1.46 NA oil immersion objective on a Zeiss LSM880 confocal equipped with an Airyscan detector. 512 × 512 pixel images were collected in 8-bit with 4-line averaging. For GFP excitation was set at 488 nm and emission at 495-550 nm; for RFP at 561 nm and 570-615 nm, respectively.

### AtLPAAT activities

The ORF for *AtLPAAT2-5* were amplified by PCR with respectively the sense/antisense primers P1539/1541 and P1543/1545 containing the XhoI and NcoI restriction sites. The PCR products were first subcloned in pGEM®-T Easy vector (Promega, Charbonnieres les Bains, France) before being cloned into the XhoI/NcoI sites of the pET-15 vector (Novagen, Merck Biosciences, Badsoden, Germany). C41 (DE3) *E. coli* bacteria (Avidis, Saint-Beauzire, France) were then transformed with the obtained plasmids. Ectopic expression of the ORF *AtLPAAT2-5* and *E. coli* membranes isolation were performed as described by Testet et al. (2005). Lysophospholipid acyltransferase reactions were carried out in 100 μl of assay mixtures (50 mM Tris-HCl, pH 8) containing 1 nmol of lysophospholipids, 1 nmol of □^14^C]oleoyl-CoA and 50 μg of membrane proteins. Reactions were incubated at 30°C and stopped at 30 min by adding 2 ml of chloroform/methanol (2:1, v/v) and 500μl of water. After the isolation of the organic phase, the aqueous phase was re-extracted with 2 ml of chloroform. The lipids were then purified by HPTLC according to Testet et al. (2005) and Ayciriex et al. (2012). The radioactivity incorporated into phospholipids was revealed using a STORM 860 PhosphorImager (GE Healthcare, Waukesha, WI) and quantified with ImageQuant TL software.

### 14-C Acetate labeling and lipid analysis

For radiolabel feeding experiments, 20 roots of 5-day-old seedlings were cut with a razor blade and placed in vials containing 2 mL MS. To start the reaction, 200 nmol (10 µCi) □1-14C] acetate (Perkin Elmer Life Sciences) were added to each vial and the reaction was stopped after 4 h with 2 ml preheated isopropanol followed by 20 min incubation at 70°C. After transfer into 6 ml glass tubes, 2ml chloroform/methanol/hydrochloric acid (100:50:1; v/v/v) were added and the mixture was incubated overnight on an orbital rotator (40 rpm) at room temperature. Subsequently, each tube was centrifuged at 1000xg for 10□min. The organic phase was collected in new tubes and the roots were re-extracted with 2 ml chloroform/methanol (2:1, v/v) for 2h. After new centrifugation, the organic phases of each sample were combined, mixed with 1.5 ml 0.9% NaCl and centrifuged at 1000g for 10 min. The organic phases were evaporated to dryness, re-suspended in 40 µl chloroform/methanol (2:1, v/v) and stored at -20□°C. Radiolabeled products were analyzed by thin-layer chromatography using HPTLC Silica Gel 60 plates (Merck) and chloroform/methanol/water/acetic acid (30:17:2:1, v/v/v/v) as solvent to separate phospholipids. They were identified by co-migration with unlabeled standards, and quantification was done by autoradiography using a Storm 860 molecular imager (GE Healthcare).

### Statistical analysis

All data analysed were unpaired (samples independent from each other). Normal distribution (Gaussian distribution) of data set was tested using Shapiro–Wilk normality test. On data normally distributed, sample homoscedasticity was assessed using Bartlett test before performing parametric tests. On data that were not normally distributed (or on data sets for which *n* <10), non-parametric tests were performed. To compare two data sets, Welch two sample *t*-test was performed on data set normally distributed, whereas Mann–Whitney test was used as non-parametric test. To compare multiple data sets, Kruskal–Wallis test was used as non-parametric test. Tukey’s test was used as a single-step multiple comparison procedure to find means significantly different from each other. All statistical tests were two-tailed (two-sided test). All statistical analyses were performed with R i386 3.1.0 software. *P*-values were as follows: ^NS^*P*-value>0.01 (not significant), \**P*<0.01, \*\**P*<0.001 and \*\*\**P*<0.0001.

## Results

### *In-vitro* AtLPAAT activities in *E*.*coli* C41 membranes

Supplementary Fig. S3 shows the alignment of the amino acid sequences of the LPAATs 2-5 from *Arabidopsis* with the sequence of the human lysophosphatidic acid acyltransferase LPAAT3 (Schmidt and Brown, 2009). The boxes indicate conserved sequences which correspond to classical motifs of the LPAAT, DHAPAT, and LPEAT acyltransferase families (Lewin et al., 1999), indicating that LPAATs 2-5 from *Arabidopsis* are clearly acyltransferases. Kim et al. (2005) found a lysophosphatidic acid acyltransferase activity for LPAT2 (AtLPAAT2) and LPAT3 (AtLPAAT3) in *E*.*coli* and yeast expressing systems but not for LPAT4 (AtLPAAT4) and LPAT5 (AtLPAAT5). Recently, Angkawijaya et al. (2019) have shown a lysophosphatidic acid acyltransferase activity for AtLPAAT5 expressed in *E*.*coli* C41 cells but not for AtLPAAT4 using this expression system. They identified AtLPAAT4 activity in *A. thaliana* transgenic plants over-expressing it. However, it is still unknown whether these enzymes only utilize LPA or whether they can also handle other lysophospholipids as substrates. To reach this goal, AtLPAAT2-5 were produced in membranes of an *E*.*coli* C41 cell line that was specifically designed for the production of membrane proteins (Miroux and Walker, 1996), and activities were measured *in vitro* as described in the experimental section and according to Testet et al. (2005). We have incubated purified membranes from *E*.*coli* C41 cells with LPA or the other lysophospholipids (lysophosphatidylcholine, lysophosphatidylethanolamine, lysophosphatidylglycerol, lysophosphatidylinositol or lysophosphatidylserine) and [^14^C]oleoyl-CoA to measure either the formation of [^14^C]PA or that of the other potentially labeled phospholipids (phosphatidylcholine, phosphatidylethanolamine, phsphatidylglycerol, phosphatidylinositol or phosphatidylserine). Controls were made with purified membranes from *E*.*coli* transformed with a pET-15B empty vector, which correspond to *E*.*coli* endogenous activities. Lysophospholipid acyltransferase activities were measured as nmol phospholipids formed/mg proteins/30min and the values of the controls were normalized to 100 for each lysophospholipid, and the corresponding activities for AtLPAAT2-5 were calculated accordingly for all the phospholipids.

Fig. 1 shows the mean activities of the four AtLPAAT2-5 from three independent experiments. As indicated, only LPA acyltransferase activities were detected for the AtLPAAT2-5 and no other lysophospholipid acyltransferase activity was observed, demonstrating that the four AtLPAATs are clearly strict lysophosphatidic acid acyltransferases. It was a very important point to determine in order to safely investigate the potential role of such enzymes and the PAs they synthesize in the functioning of the secretory pathway.

**Fig.1.**
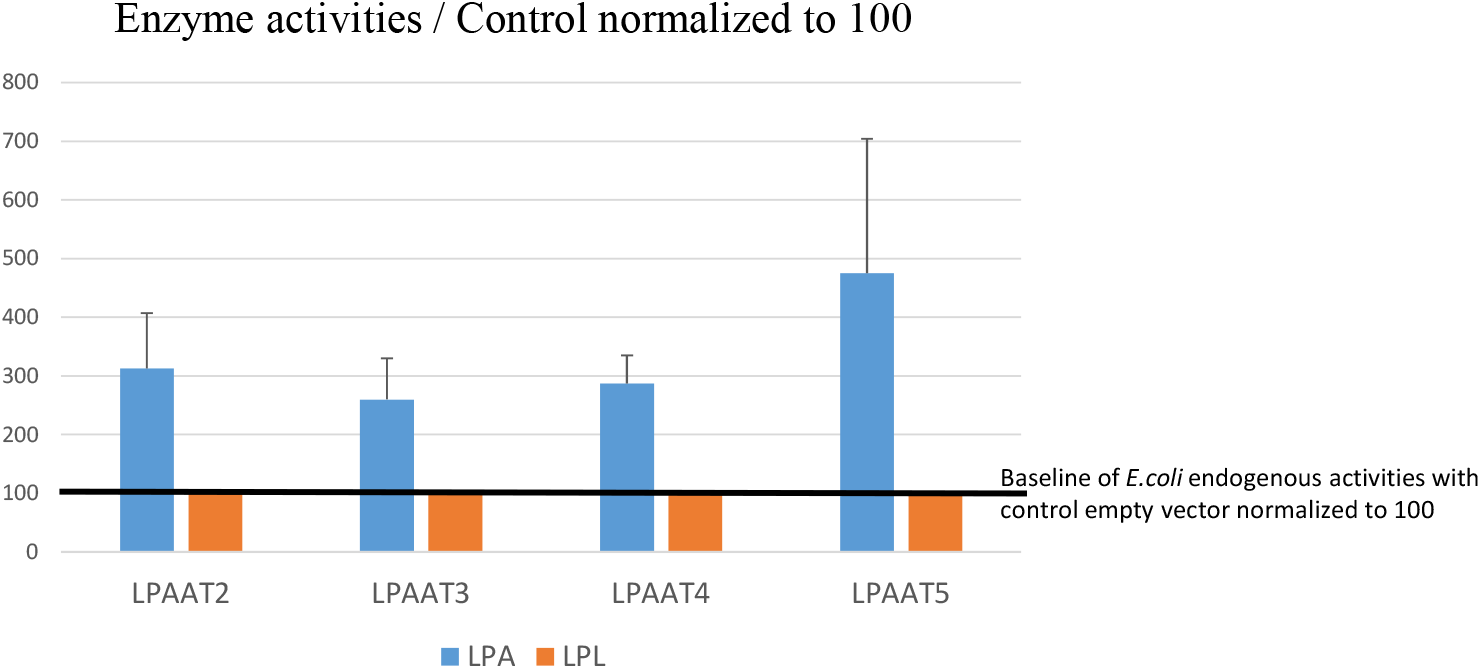
*In-vitro* AtLPAAT2-5 LPA acyltransferase activities in *E*.*coli* C41 membranes. *AtLPAAT2-5* were produced in membranes of *E. coli* C41 cell line that was specifically designed for the production of membrane proteins (Miroux and Walker, 1996), and activities were measured *in vitro* as described in the experimental section and according to Testet et al. (2005). Activities were tested with lysophosphatidic acid (LPA) and the other lysophospholipids (LPL: either lysophosphatidylcholine, lysophosphatidylethanolamine, lysophsphatidylglycerol, lysophosphatidylinositol or lysophosphatidylserine). Lysophospholipid acyltransferase activities were measured as nmol phospholipids formed/mg proteins/30min and the values of the controls (corresponding to purified membranes from *E*.*coli* transformed with pET-15B empty vector) were normalized to 100 for each lysophospholipid and the corresponding activities for AtLPAAT2-5 were calculated accordingly for all the phospholipids. Only LPA acyltransferase activities were detected for the AtLPAAT2-5 and no other lysophospholipid acyltransferase activity was detected, determining that AtLPAAT2-5 are strict LPA acyltransferases.

The levels of activities of PA synthesis were about 1.25 nmol/mg/30 min for AtLPAAT2, 1.08 nmol/mg/30 min of PA for AtLPAAT3, 1.72 nmol/mg/30 min of PA for AtLPAAT4 and 1.75 nmol/mg/30 min of PA for AtLPAAT5, showing similar levels of activities in *E. coli* C41 cell membranes, which does not necessarily reflect their level of activities *in planta*.

### Subcellular localization of AtLPAATs in *Arabidopsis*

The AtLPAAT2 has been shown to be located in the ER (Kim et al., 2005), and very recently, Angkawijaya et al. (2019) have shown that AtLPAAT4 and AtLPAAT5 are located in the ER. However, the subcellular localization of the AtLPAAT3 has not been determined yet. To investigate its membrane localization, roots of an *Arabidopsis* stable line (5 days after germination) expressing both GFP-LPAAT3 and the ER marker mCherry-HDEL were observed. GFP-LPAAT3 was not diffusely localized in the ER network as the other AtLPAATs but as dots very close to the ER (Fig. 2A-F). We further investigated AtLPAAT3 localization using an heterologous expression system in tobacco leaf epidermal cells in which the ER is more dynamic and accessible to look at ER domains than in root cells. Interestingly, a transient expression in this system of tagRFP-LPAAT3 and the ER-export sites (ERES) marker SAR1-GFP showed a colocalization (Fig. 2G-I) which indicates that the AtLPAAT3 dots observed in *Arabidopsis* roots may correspond to ERES at least for some of them. Looking at the AtLPAAT3 sequence, we found the di-acidic motifs (D74-X-E76 and D293-X-E295) which could serve as ER export signals and may explain the different localization compared to the other LPAATs. Transient expressions in *Arabidopsis* cotyledons of GFP-LPAAT3 di-acidic mutants 1 (D74G/A75/E76G) and 2 (D293G/L294A/E295G) with the ER marker mCherry-HDEL were realized (Fig. 3). In both cases, the mutated GFP-LPAAT3 was significantly redistributed into the ER membranes network, indicating that these motifs may be functional. However, such export motifs are usually active to conduct the efficient export of proteins outside of the ER to reach subsequent compartments in the secretory pathway but GFP-LPAAT3 was not observed in compartments distant from the ER. Altogether our data suggest that AtLPAAT3 is localized in ER domains near the ER-Golgi interface rather than distributed into ER membranes as found for AtLPAAT2, AtLPAAT4 and AtLPAAT5.

**Fig.2.**
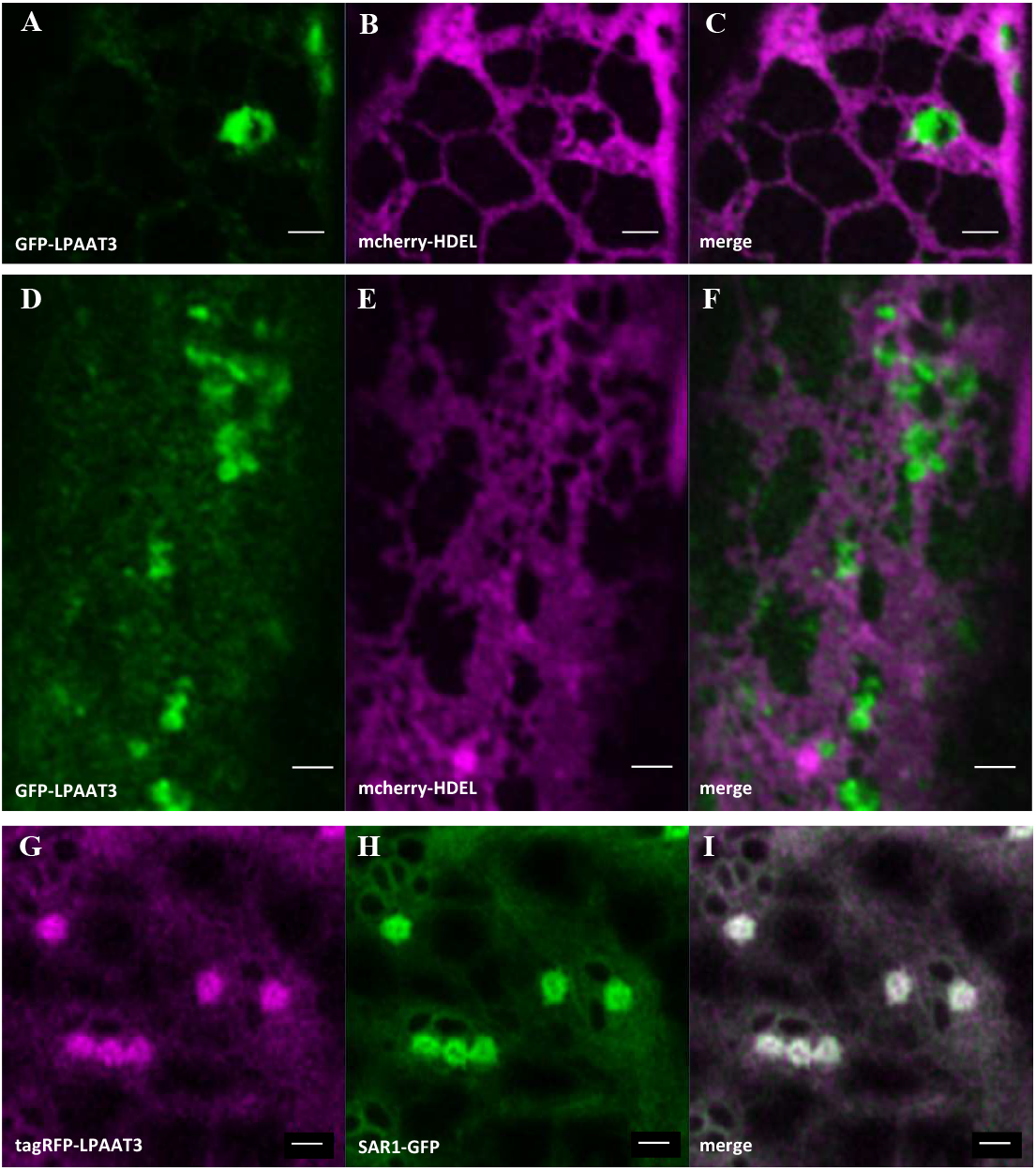
Subcellular localization of AtLPAAT3 in various plant models. Roots, 5 days after germination, of an *Arabidopsis* stable line expressing both GFP-LPAAT3 (A,D) and the ER marker mCherry-HDEL (B,E). AtLPAAT3 was found in punctuated structures very close to the ER (C,F). A transient expression of tagRFP-LPAAT3 (G) and the ERES marker SAR1-GFP (H) in *Nicotiana tabacum* leaf epidermis cells suggested that the punctuated structures observed for AtLPAAT3 in roots could potentially correspond to ERES (I). Scale bars 1µm.

**Fig.3.**
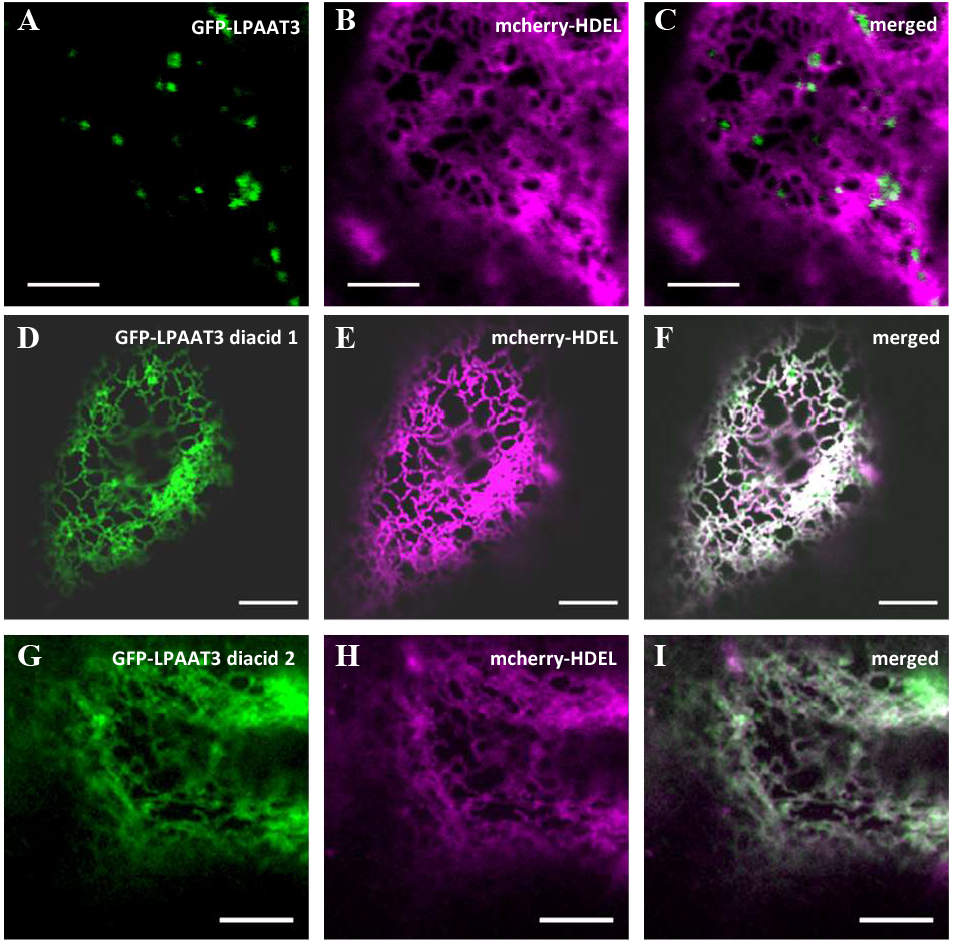
Subcellular localization of AtLPAAT3 depends on active ER export motifs. Transient expression in *Arabidopsis* cotyledons of LPAAT3 GFP fused native forms (A), GFP-LPAAT3 di-acidic mutant 1 (D), GFP-LPAAT3 di-acidic mutant 2 (G) and the ER marker mCherry-HDEL (B,A,H). The mutation of each diacidic motif induces redistribution of LPAAT3 into ER membranes network (F,I). Scale bars 5µm.

### AtLPAAT2, AtLPAAT4 and AtLPAAT5 do not cycle between the ER and the Golgi

As shown in Supplementary Fig. S3, AtLPAAT2, AtLPAAT4 and AtLPAAT5, but not AtLPAAT3, show potential di-lysine motifs at their C-termini (respectively KXK, KXKXX and KXXXK motifs) which may indicate that these enzymes could eventually cycle between the ER and Golgi/post-ER compartments. To investigate this possibility, we mutated the potential cycling motifs (GFP-LPAAT2 *K387A/K389A*, GFP-LPAAT4 *K374A/K376A* and GFP-LPAAT5 *K371A/K375A*) and looked at the expression of these mutant fluorescent constructs in *Arabidopsis* cotyledons together with the ER marker mCherry-HDEL. As shown in Fig. 4, all the mutated versions of the AtLPAAT2, AtLPAAT4, and AtLPAAT5 were still located in the ER and did not show any fluorescent dots which could correspond to Golgi/post-ER compartments. Hence these enzymes do not cycle between the ER and the Golgi or post-ER compartments, excluding any role of these proteins at the level of the Golgi membranes but only at the level of the ER membranes.

**Fig.4.**
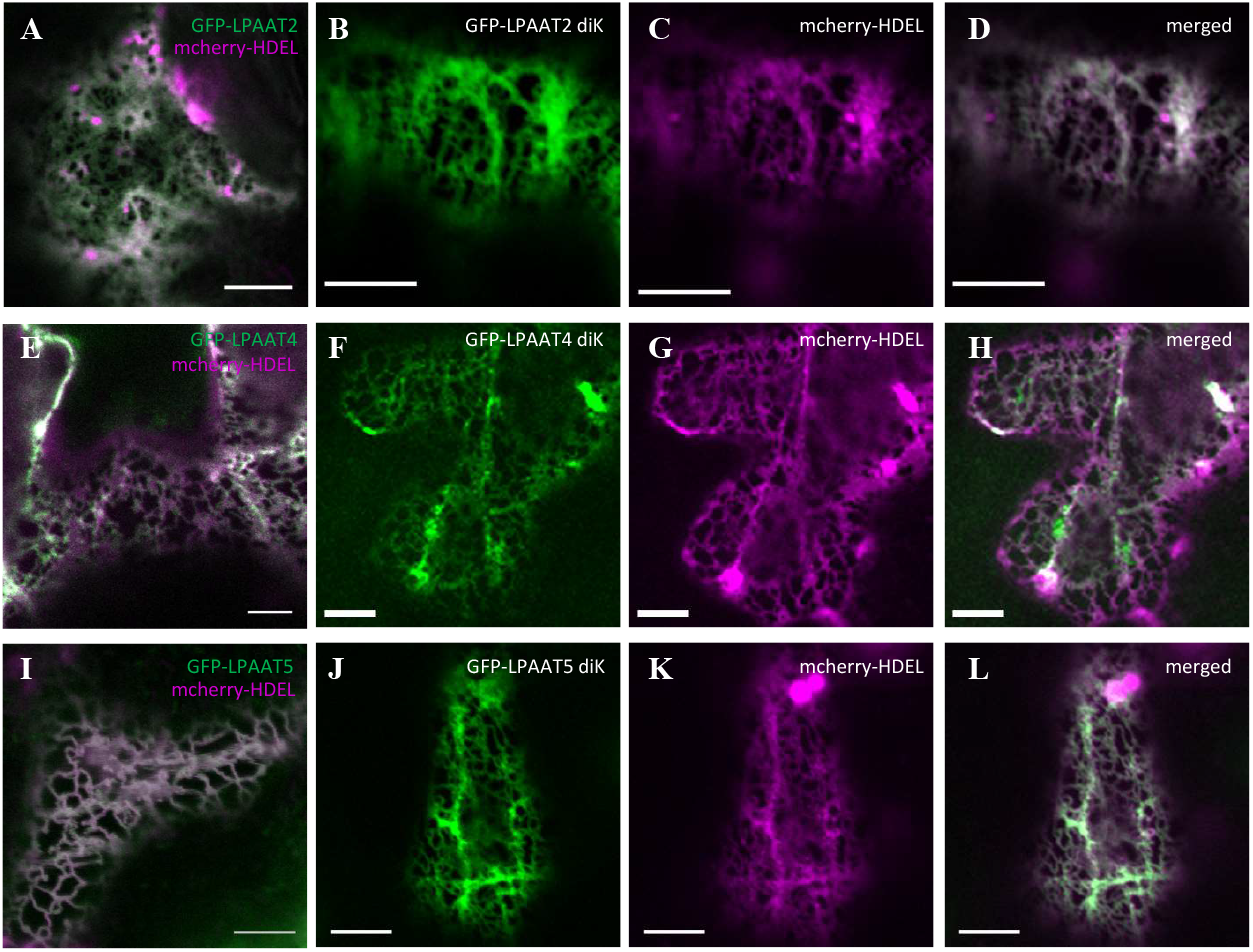
AtLPAAT2, AtLPAAT4 and AtLPAAT5 do not cycle between the ER and the Golgi. Transient expression in *Arabidopsis* cotyledons of GFP-LPAAT2 diK mutant (B), GFP-LPAAT4 diK mutant (F), GFP-LPAAT5 diK mutant (J) and the ER marker mCherry-HDEL (C,G,K). Results are compared to transient co-expression of GFP fused native forms of AtLPAAT2 (A), AtLPAAT4 (E) and AtLPAAT5 (I) with the ER marker mCherry-HDEL. Mutations of AtLPAAT2, AtLPAAT4 and AtLPAAT5 diK motifs do not impact their location to the ER membranes network (D,H,L). Scale bars 5µm.

AtLPAAT2 being ubiquitous and likely the main source of PA for the *de novo* synthesis of phospholipids in the ER (Kim et al., 2005; Angkawijaya et al., 2017), we may consider that some of the other ER AtLPAATs may be involved in the trafficking machinery from the ER. Therefore, we decided to focus on the three other AtLPAATs to investigate their potential requirement in the secretory pathway. Since AtLPAAT3-5 are similarly expressed in roots (Supplementary Fig. S4), we chose primary root growth as a phenotypic readout to look at the functioning of the secretory pathway.

### Primary root growth phenotype and sensitivity to CI976 of *lpaat* mutants

Since an *AtLPAAT2* KO mutant is lethal (Kim et al., 2005), we first looked at the primary root growth of the single KO mutants *lpaat3-1* (Angkawijaya et al., 2017), *lpaat4-2 and lpaat5-2* (different alleles from those used by Angkawijaya et al., 2019) five days after germination. The caracterisation of *lpaat* insertion mutant lines and T-DNA positions is shown in Supplementary Fig. S1. As shown in Fig. 5A, none of these mutants had a decrease in their primary root growth which could be due to complementation by each other. Therefore, we produced the three double mutants *lpaat3-1;lpaat4-2, lpaat3-1;lpaat5-2 and lpaat4-2;lpaat5-2* and looked at the primary root length five days after germination (Fig. 5B). A significant but weak root length phenotype was observed for all the double mutants. As a consequence, we decided to produce the triple mutant *lpaat3-1;lpaat4-2;lpaat5-2* and looked at again the primary root length of these mutants. Surprisingly, primary growth was not inhibited but stimulated in the triple mutant (Supplementary Fig. S5). Since the over-expression of *AtLPAAT2* stimulates the *de novo* synthesis of phospholipids, resulting in an enhanced primary root growth in phosphate-starved *Arabidopsis* seedlings (Angkawijaya et al., 2017), we wondered whether the expression of *AtLPAAT2* was boosted in the triple mutant *lpaat3-1;lpaat4-2;lpaat5-2*. To check this hypothesis, semi-quantitative RT-PCR and real-time RT-PCR analyses of *AtLPAAT2* transcripts were performed in roots five days after germination (Supplementary Fig. S6). As shown, the expression of *AtLPAAT2* was increased in the triple mutant *lpaat3-1;lpaat4-2;lpaat5-2* in comparison to the wild-type but not in the double mutant *lpaat4-2;lpaat5-2*. This result shows that, in the context of the triple mutant *lpaat3-1;lpaat4-2;lpaat5-2, AtLPAAT2* is over-expressed and could compensate the absence of the three other AtLPAATs and therefore resulting in an increase in primary root growth. In agreement, the triple mutant showed the same level of PA amounts than the WT (see below Fig.7A).

**Fig.5.**
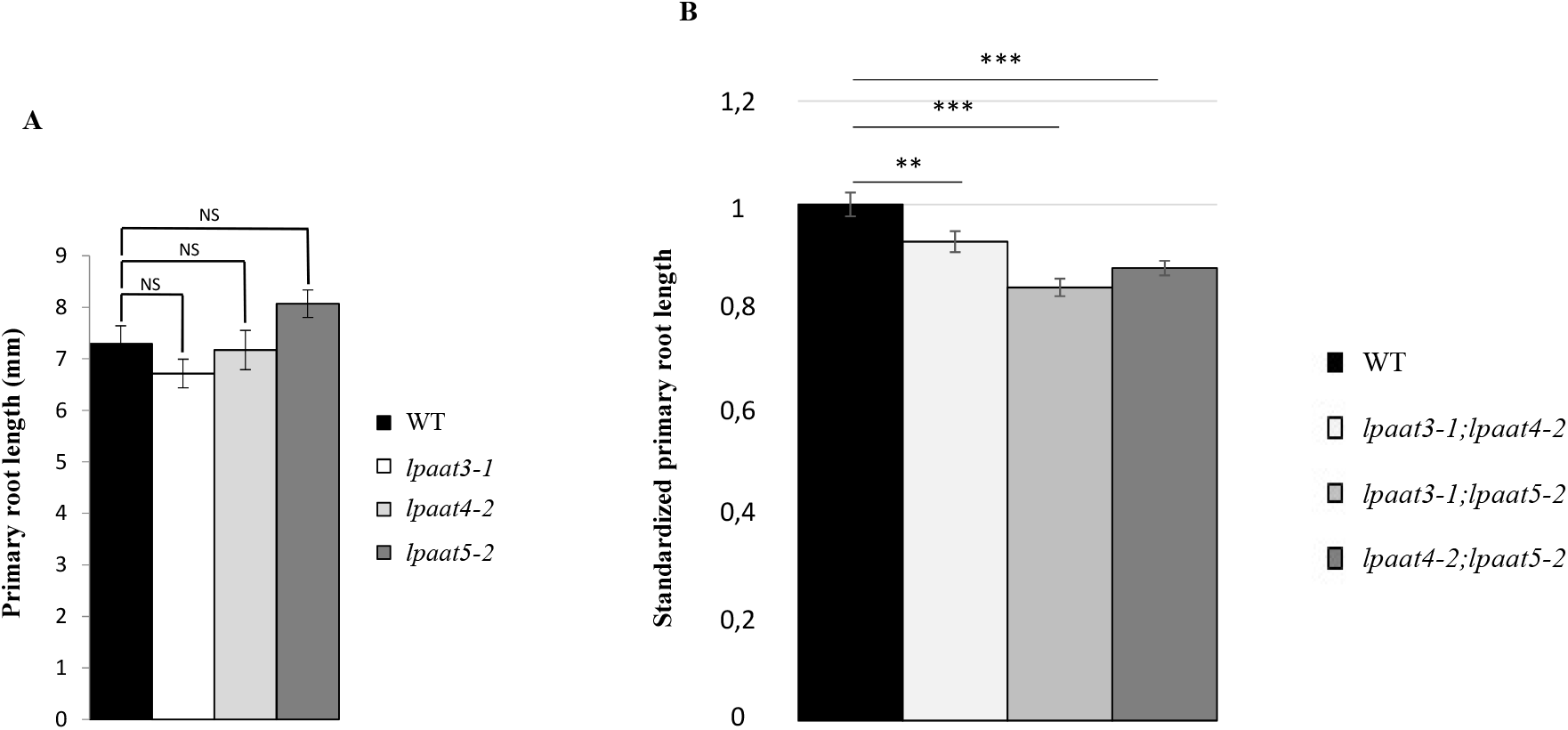
Primary root growth of *lpaat* KO mutants (A) is not affected and that of *lpaat* KO double mutants (B) is significantly but only slightly affected. Primary root length was measured 5 days after germination. Data are mean values ± SE, n = 40 (*lpaat* KO mutants), n = 80 (*lpaat* KO double mutants). Statistics were done by Kruskal-Wallis rank sum test, NS = not significant, **P-value <0.01, ***P-value < 0.001.

Therefore, how can we determine the putative role of AtLPAATs in the secretion pathway with double mutants showing only a weak growth-inhibition phenotype and a triple mutant showing even an increase in root growth? To solve this puzzling question, we took advantage of our experience in combining genetic and biochemical approaches as we did before to investigate the role of sphingolipids in protein trafficking at the Golgi complex (Melser et al., 2010; Wattelet-Boyer et al., 2016). CI976 being an inhibitor of LPAT activities that interferes with both COPII- and COPI-dependent membrane traffic processes (Brown et al., 2008; Schmidt and Brown, 2009; Yang et al., 2011), we decided to look at the effects of this inhibitor on primary root growth in the double mutant *lpaat4-2;lpaat5-2* (with no increase in the expression of AtLPAAT2) and the triple mutant *lpaat3;lpaat4;lpaat5* compared to the WT. The choice of the double mutant *lpaat4-2;lpaat5-2* was also based on the fact that both AtLPAAT4 and AtLPAAT5 have the same diffuse localization in the ER and to focus the inhibitor on the AtLPAAT3 which may be at least partially associated with the ERES. While a small decrease in primary root growth (but not statistically significant) was observed in WT and no effect at all in the triple mutant *lpaat3-1;lpaat4-2;lpaat5-2* (as expected due to overexpression of *AtLPAAT2)*, a significant decrease in growth was observed in the double mutant *lpaat4-2;lpaat5-2* treated with 10µM of CI976 as shown in Fig. 6. Higher concentrations of CI976 induced a decrease in primary root growth in all the lines, justifying 10µM as the functional concentration of the inhibitor CI976 to be used to perform our investigations on primary root growth and subsequently on protein transport. We also checked whether the inhibitor could enhance the expression of *AtLPAAT2* and as shown in the Supplementary Fig. S6, this expression was similarly increased in both the WT and the double mutant *lpaat4-2;lpaat5-2* but to a much lesser extent than in the triple mutant *lpaat3-1;lpaat4-2;lpaat5-2*. Therefore, treating the double mutant *lpaat4-2;lpaat5-2* with 10µM of CI976 permitted to establish primary root growth phenotypic conditions and allowed investigating the efficiency of protein trafficking (see below).

**Fig.6.**
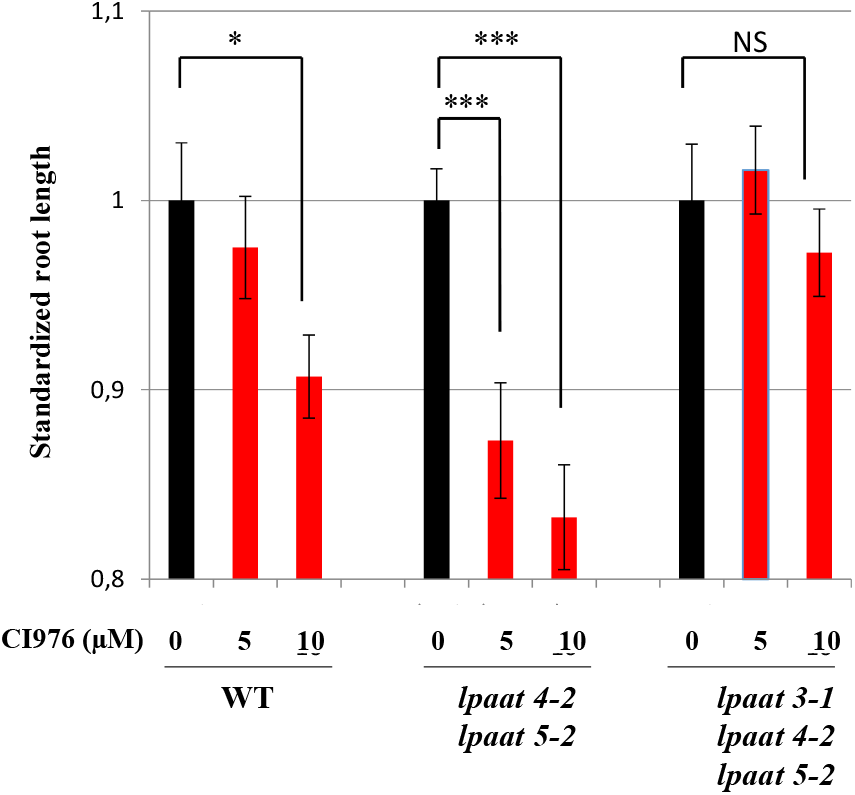
Sensitivity of WT, double mutant *lpaat4-2;lpaat5-2* and triple mutant *lpaat3-1;lpaat4-2;lpaat5-2* lines to CI976 treatment. Seedlings were grown on MS agar medium plates containing different concentrations of CI976 (0, 5, 10µM). Primary root length was measured 5 days after germination and standardized to the untreated condition for each line. Data are mean values ± SE from three biological replicates, n = 60. The asterisks indicate significant difference between untreated condition (black bar) and CI976 treated conditions (red bars). Statistics were done by non-parametric Kruskal-Wallis rank sum test, \*\*\**P*-value < 0.0001, NS = not significant.

### PA biosynthesis in *Arabidopsis* roots is only affected in the double mutant treated by CI976

Since the perturbation of the AtLPAAT activities led to a decrease in primary root growth, we determined the amount of neo-synthesized PA in *Arabidopsis* roots of the WT, the double mutant *lpaat4-2;lpaat5-2* and the triple mutant *lpaat3-1;lpaat4-2;lpaat5-2* with or without CI976 treatment. As shown in Fig. 7A, the amount of neo-synthesized PA in the untreated double mutant *lpaat4-2;lpaat5-2* was reduced to 60% of that of the untreated WT and triple mutant *lpaat3-1;lpaat4-2;lpaat5-2*, this was correlated only to a weak primary root growth phenotype as mentioned before (Fig. 5). Treatment of the WT and the triple mutant *lpaat3-1;lpaat4-2;lpaat5-2* with the CI976 led to a level of neo-synthesized PA similar to that found in the untreated double mutant *lpaat4-2;lpaat5-2* (Fig. 7A). Treating the double mutant *lpaat4;lpaat5* with CI976 led to an additional decrease in PA which reached only about 30-35% of that of the untreated WT and triple mutant *lpaat3-1;lpaat4-2;lpaat5-2* (Fig. 7A). In these conditions, we observed a higher primary root growth phenotype (Fig. 6). This may indicate a concentration threshold for PA with no clear or very weak phenotype at concentrations above this level and a clear primary root growth phenotype at concentrations below (Fig. 6).

**Fig.7.**
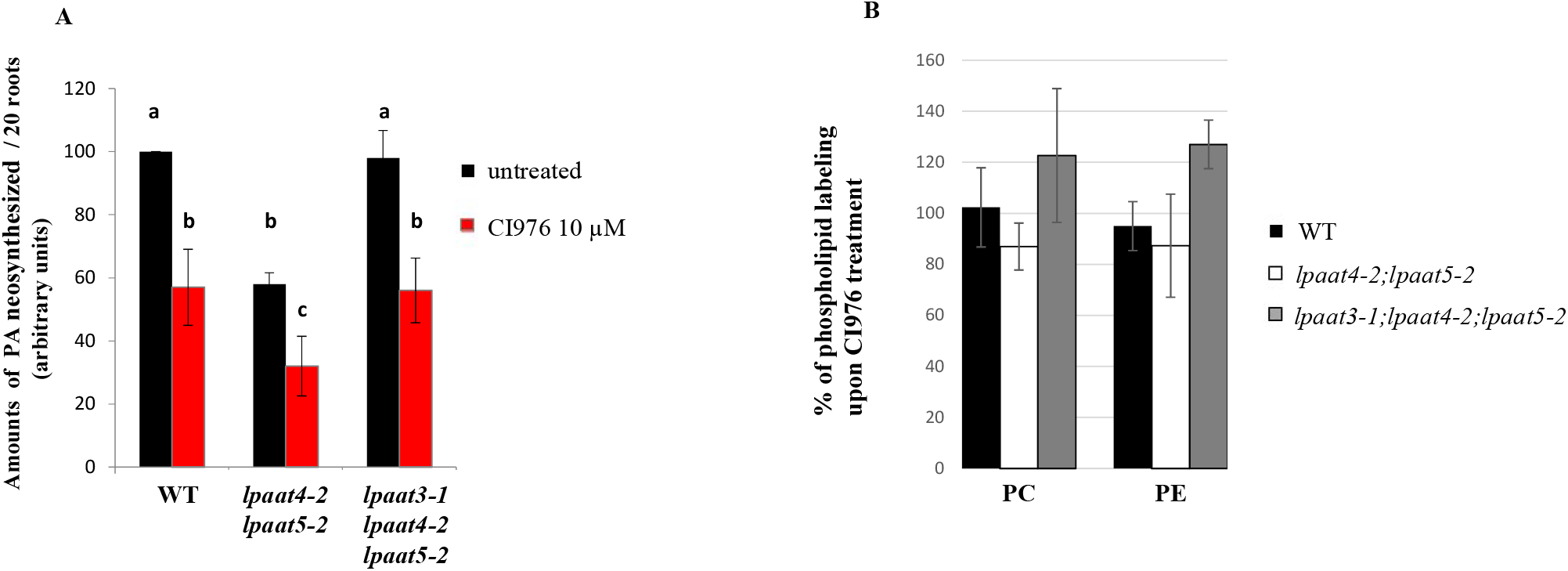
CI976 treatment affects the neo-synthesis of PA but not that of the major phospholipids in Arabidopsis roots. Seedlings were grown on MS agar medium plates or MS agar medium plates complemented with 10 μM CI976 or not. Primary roots from 20 seedlings were collected 5 days after germination for both conditions and incubated 4 hours with [14C] acetate +/- 10 µMCI976. Lipids were extracted and separated by HPTLC. A. Amounts of neo-synthesized [14C] PA, [14C] PC and [14C] PE were quantified for the WT line and the mutant lines *lpaat4-2;lpaat5-2* and *lpaat3-1;lpaat4-2;lpaat5-2*. The amounts of [14C] PA, produced were calculated for each line without treatment (black bar) or under treatment (red bar) by taking as 100 the amounts of [14C] PA produced in the WT line without treatment. Data are mean values ± SD from three biological replicates. Statistics were made with the non-parametric Kruskal-Wallis test, similar letters above bars indicate that dataset are not significantly different, b/a and c/b: P-values < 0.0001. B. [14C] PC and [14C] PE produced were quantified in the WT line, the double mutant line *lpaat4-2;lpaat5-2* and the triple mutant line *lpaat3-1;lpaat4-2;lpaat5-2* upon CI976 treatment, and compared to the amounts measured for each line without treatment. The % of labeling of PC and PE in the presence of CI976 was expressed as compared to the untreated conditions taken as equal to 100. Data are mean values ± SD from three biological replicates. Statistics were made with the non-parametric Kruskal-Wallis test and the p values for PC (0.393) and PE (0.288) show no significant differences.

A small decrease in the amount of phospholipids was observed in the untreated double mutant *lpat4-1;lpat5-1* in the study of Angkawijaya et al. (2019). Therefore, an additional question was to determine whether the primary root growth phenotype observed in the CI976-treated double mutant *lpaat4-2;lpaat5-2* was only due to a decrease in the neo-synthesis of PA or whether the neo-synthesis of the major phospholipids was also affected by the CI976 treatment and could contribute to the primary root growth phenotype. For this, we measured the level of the neo-synthesis of the two major phospholipids phosphatidylcholine (PC) and phosphatidylethanolamine (PE) from [^14^C] acetate in *Arabidopsis* roots of the WT, the double mutant *lpaat4-2;lpaat5-2* and the triple mutant *lpaat3-1;lpaat4-2;lpaat5-2* treated with CI976. Beside the significant decrease in labeled PA in the CI976-treated double mutant *lpaat4-2;lpaat5-2* compared to the WT and the triple mutant *lpaat3-1;lpaat4-2;lpaat5-2* (Fig. 7A), we did not observe any significant decrease in the amounts of labeled PC and PE (Fig. 7B), indicating that the synthesis capability of these major phospholipids were similar in the CI976-treated double mutant *lpaat4-2;lpaat5-2* compared to the WT and the triple mutant *lpaat3-1;lpaat4-2;lpaat5-2*. Therefore, the primary root growth phenotype observed in the CI976-treated double mutant *lpaat4-2;lpaat5-2* can be mainly attributed to the disturbance of PA metabolism unrelated to the neo-synthesis of the major phospholipids. Therefore, by combining genetic and pharmacological approaches, we determined the best conditions to investigate the potential role of these LPAATs in the functioning of the secretory pathway.

### Efficiency of protein secretion in the CI976 treated double mutant *lpaat4-2;lpaat5-2*

To investigate the efficiency of protein trafficking in the CI976 treated double mutant *lpaat4-2;lpaat5-2*, we decided to look *in situ* at several plasma membrane markers (H+ATPases, PIN2 and PIP2,7) already characterized in their trafficking to the plasma membrane (Melser et al., 2010; Wattelet-Boyer et al., 2016; Hachez et al., 2014).

We first investigated the transport to the plasma membrane of H+ATPases by an immunocytochemistry approach. Supplementary Fig. S7 shows that the transport to the plasma membrane of H+ATPases was not affected in the CI976-treated double mutant *lpaat4-2;lpaat5-2* compared to the WT. As a consequence, the transport to the plasma membrane of H+ATPases did not seem to require the LPAAT-dependent production of PA.

To investigate the impact of LPAATs on PIN2 transport to the plasma membrane, we crossed the stable line pPIN2::PIN2–GFP (Xu and Scheres, 2005) with the WT and the double mutant *lpaat4-2;lpaat5-2* line and treated or not the growing roots with 10µM CI976. As shown in Fig. 8, we observed an increase in intracellular PIN2, indicating that the transport of PIN2 was disturbed. As an approach to try identifying the compartment(s) where PIN2 was retained, we used an immunostaining strategy using antibodies raised against various compartments of the secretory pathway, Echidna (ECH, marker of the SYP61 compartment, Gendre et al., 2011 and Boutté et al., 2013), SAR1 (an ERES marker, Hanton et al., 2007) and Membrine11 (Memb11, a cis-Golgi marker, Marais et al., 2015). We performed immunostaining upon CI976 treatment of the double mutant *lpaat4-2;lpaat5-2* expressing PIN2-GFP (Supplementary Fig. S8A-J). A significant co-localization was observed with ECH, but no significant co-localization was observed with SAR1 and Memb11 (Supplemental Figure S8K), indicating that PIN2 was mainly retained at the level of the TGN.

**Fig.8.**
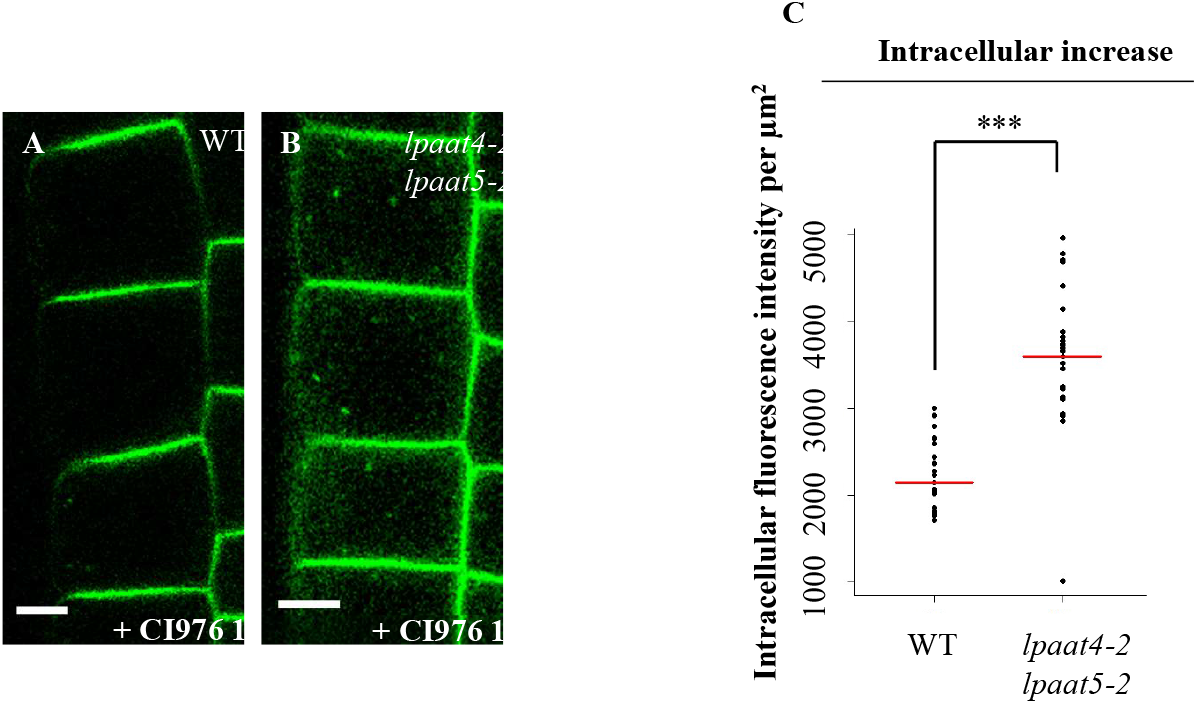
Auxin carrier PIN2-GFP trafficking is altered in double mutant *lpaat4-2;lpaat5-2* upon CI976 treatment. Localization of PIN2-GFP in WT (A) and *lpaat4-2,lpaat5-2* (B) background upon 10μM CI976 treatment. Sum of fluorescense intensity per μm2 was calculated in the cytoplasm for each line upon treatment. The intracellular increase in PIN2-GFP in *lpaat4-2,lpaat5-2* upon CI976 treatment is shown (C) (n=27 cells quantified over 8 independent roots). All the data were represented for each line (black dots) with the median of each dataset (red bar). Statistics were done by non-parametric Kruskal-Wallis rank sum test, ***P-value < 0.0001.

To investigate the impact of LPAATs on PIP2;7 transport to the plasma membrane, we looked at the subcellular location of PIP2;7 in the WT, the double mutant *lpaat4;lpaat5* and the triple mutant *lpaat3-1;lpaat4-2;lpaat5-2* treated with 10µM CI976. For this purpose, an immunocytochemistry approach to reveal PIP2;7 *in situ* localization was used. A whole-mount immunolabelling of *Arabidopsis* roots was performed as described previously (Boutté and Grebe, 2014). As shown in Fig. 9, a decrease in the mean fluorescence ratio of plasma membrane to cytoplasm was observed for the double mutant *lpaat4-2;lpaat5-2* compared to the WT and the triple mutant *lpaat3-1;lpaat4-2;lpaat5-2* which corresponded to both a decrease in PIP2;7 in the plasma membrane and an increase in the protein amount in the cytoplasm. It was reported that PIP2;7 is highly regulated under salt stress (Pou et al., 2016) which also increases the *AtLPAAT4* gene expression in roots (Supplementary Fig. S9). We first checked for a sensitivity of the double mutant *lpaat4-2;lpaat5-2* to salt stress. The double mutant *lpaat4-2;lpaat5-2* was effectively more sensitive to salt stress than the WT and the triple mutant *lpaat3-1;lpaat4-2;lpaat5-2* at 50mM NaCl (Fig. 10). Higher salt concentrations up to 150mM gave stronger phenotypes but without significant differences between the WT and the mutant lines. Interestingly, when we looked at the sensitivity of the different lines to salt stress in the presence of CI976 (Fig. 10), we observed that the double mutant *lpaat4-2;lpaat5-2* became less sensitive in the presence of the drug, and reached the value observed with the WT and the triple mutant *lpaat3-1;lpaat4-2;lpaat5-2* (Fig. 10). These results can be easily explained by the fact that PIP2;7 localized less to the plasma membrane of double mutant *lpaat4-2;lpaat5-2* root cells under CI976 treatment than in WT and triple mutant *lpaat3-1;lpaat4-2;lpaat5-2* (Fig. 9). By complementing the double mutant *lpaat4-2;lpaat5-2* by overexpression of *AtLPAAT4* (*AtLPAAT4* relative transcript abundance in the double mutant *lpaat4-2;lpaat5-2* is shown in the Supplementary Fig. S10), we observed a partial restoration of the localization of PIP2;7 at the plasma membrane (Fig.9). These results associated with the fact that salt stress increases *AtLPAAT4* gene expression in roots (Pou et al., 2016) suggest that PA formed by some LPAATs is involved in both the correct functioning of the secretory pathway and lipid signaling processes.

**Fig.9.**
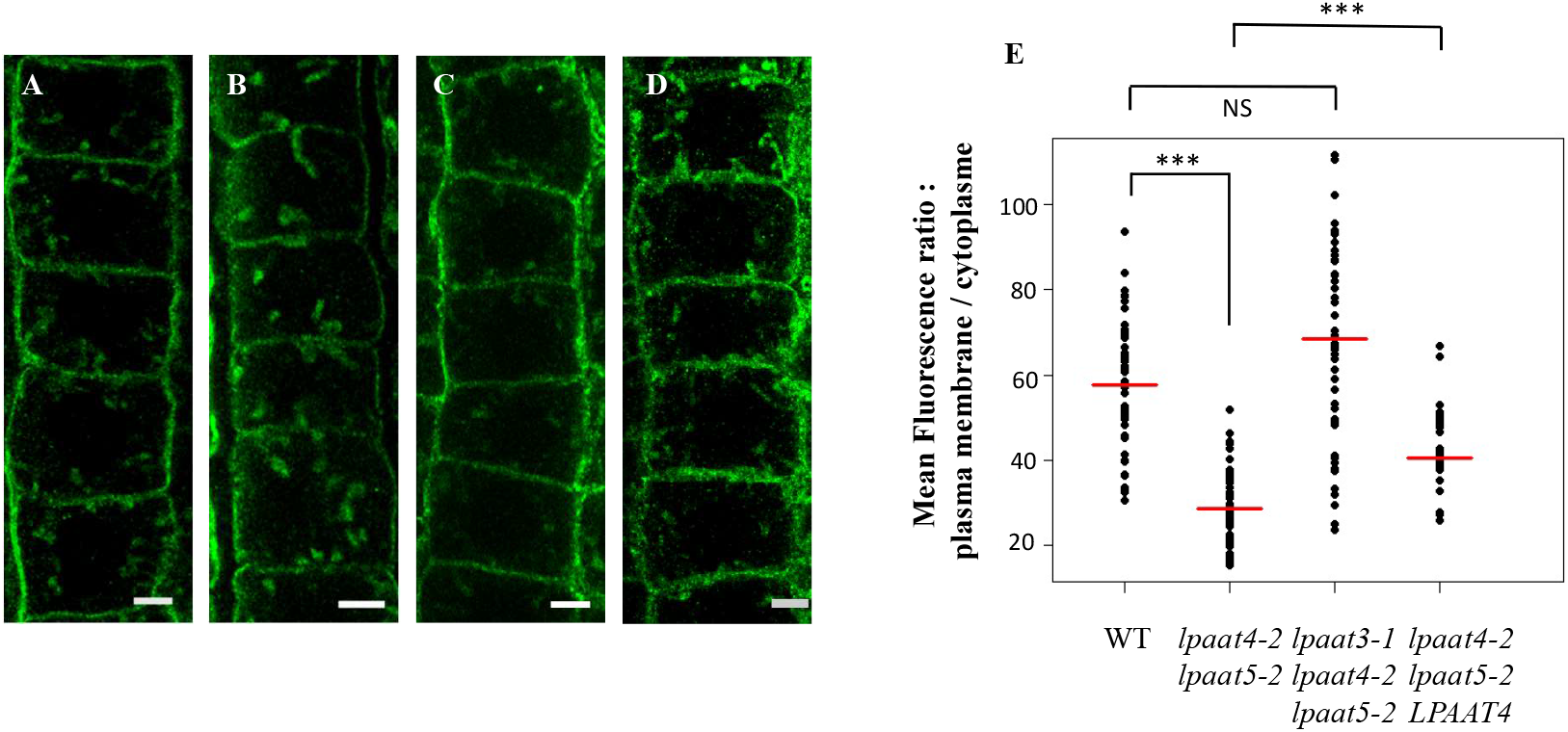
Aquaporin PIP2;7 trafficking is altered in double mutant *lpaat4-2;lpaat5-2* upon CI976 treatment. Immunolocalization of the Aquaporin PIP2;7 in root epithelial cells from *Arabidopsis* wild-type (A), double mutant *lpaat4-2,lpaat5-2* (B), *triple mutant lpaat3-1;lpaat4-2,lpaa5-2* (C) and double mutant *lpaat4-2,lpaat5-2* over-expressing LPAAT4 (D) lines upon CI976 treatment. Scale bar represents 5μm. Mean fluorescence intensity was measured at the plasma membrane and in the cytosol for each line upon treatment. The ratio of fluorescence intensity between the plasma membrane and the cytosol was calculated. (E) All the ratios were represented for each line (black dots) with the median of each dataset (red bar). Statistics were done by non-parametric Kruskal-Wallis rank sum test, ***P-value <0.0001, NS: not significant.

**Fig 10.**
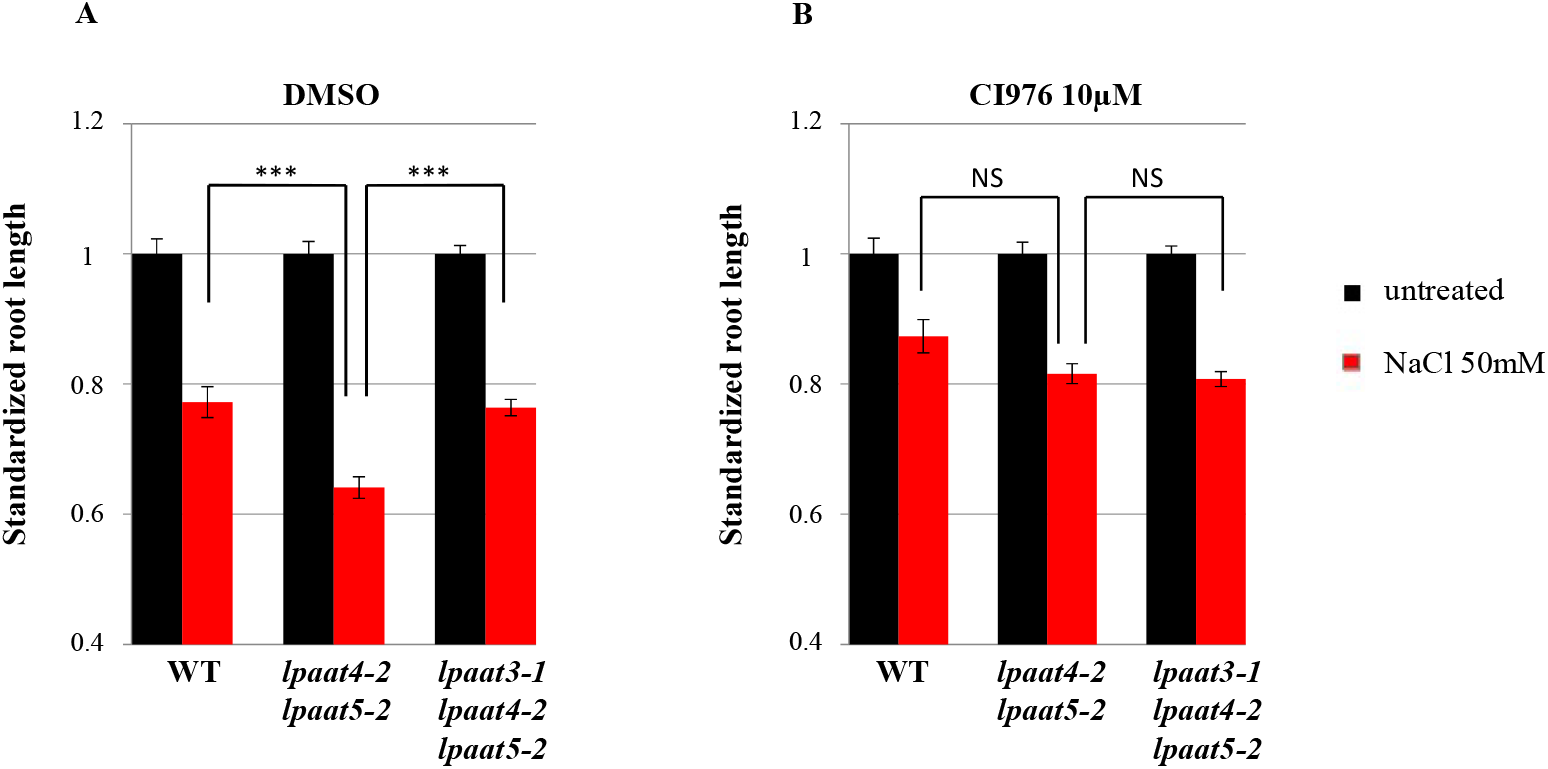
Double mutant *lpaat4-2;lpaat5-2* is highly sensitive to salt stress as compared to WT, and triple mutant *lpaat3-1;lpaat4-2;lpaat5-2*, effect on primary root length. Seedlings were grown on MS agar medium plates supplemented or not with NaCl 50 mM. The same experiment was performed with (A) or without (B) CI976 treatment. Primary root length were measured 5 days after germination. Values were standardized to WT for each condition. Results indicate a higher sensitivity from the double mutant *lpaat4-2;lpaat5-2* to salt stress in comparison to the wild-type and triple mutant *lpaat3-1;lpaat4-2;lpaat5-2* lines (C), while this sensitivity is lost upon CI976 treatment (B). Data are mean values ± SE from three biological replicates (n = 150). Statistics were done by non-parametric Kruskal-Wallis rank sum test, \*\*\**P*-value < 0.0001.

The increase in the amounts of PIP2;7 and PIN2 in intracellular membranes was interpreted as a consequence of a decrease in the transport of these proteins to the plasma membrane. However, the same result could have been the consequence of an increase in their internalization. To test whether this might have been the case, we measured endocytosis of the marker FM4-64 in both WT and double mutant *lpaat4-2;lpaat5-2* lines treated with CI976. As shown in the Supplementary Fig. S11, it was found that not only the internalization of the marker FM4-64 was not increased but even a slight decrease was observed. As a consequence, it is unlikely that the increase of PIP2;7 and PIN2 was due to an increase in endocytosis. Therefore, our results strongly argue that a decrease in the transport efficiency of PIP2;7 and PIN2 to the plasma membrane explains the increase of these protein amounts in intracellular membranes.

In addition, our results show that the disruption of the AtLPAAT activities affected to some extent the efficiency of the secretory pathway followed by PIP2;7 and PIN2 but not H+ATPases, suggesting either different sensitivities to PA of their trafficking process or different requirements for it.

## Discussion

PA is a central phospholipid metabolic intermediate which is essential for the *de novo* synthesis of membrane lipids and also a key second messenger and source of other signalling lipids for numerous signalling pathways activated during diverse stress conditions (Pokotylo et al., 2018; Yao and Xue, 2018). PA, like phosphoinositides and phosphatidylserine, is involved in the differentiation of various electrostatic compartments in the cell (Platre et al., 2018) and has been shown to interact more or less specifically with numerous proteins engaged in a large variety of cell functions (Pokotylo et al., 2018). Beside these critical functions, PA may also contribute to the function of the plant secretory pathway through its physico-chemical properties (Furt and Moreau, 2009; Boutté and Moreau, 2014). From a mechanical point of view, PA is a cone-shaped lipid with the property to favour negative membrane curvature, and its precursor lysophosphatidic acid has the tendency to favor positive membrane curvature due to its inverted cone shape. Both of these lipids can stimulate physico-chemical mechanisms linked to membrane morphodynamics according to the leaflet of the membrane where they are produced (Boutté and Moreau, 2014). Yang et al. (2011) have remarkably demonstrated the interplay between phospholipase A2 and LPAAT in regulating COPI vesicles versus tubules formation from Golgi membranes in mammalian reconstituted systems. More recently, lysophospholipids have been shown to be critical in the formation of COPII vesicles by inducing the required membrane deformation (Melero et al., 2018), and they are involved in PIN intracellular trafficking (Lee et al., 2010) and pollen germination and development (Kim et al., 2011). In addition to enzymes such as phospholipases which have been shown to be involved in membrane trafficking in plant cells (Li et al., 2007; Lee et al., 2010; Li et al., 2011; Kim et al., 2011), acyltransferases which are potentially involved in lipid metabolism in the Land’s cycle may have an impact on membrane morphodynamics in plant cells (Boutté and Moreau, 2014).

Interestingly, Pleskot et al. (2012) have shown different roles of PA produced by either phospholipases D or diacylglycerol kinases in pollen tube growth. Their results strongly suggested that several pools of PA may exist according to the biosynthetic pathway followed by PA and the cellular process concerned. We may also consider that different LPAATs could produce different pools of PA linked to various cellular processes (*de novo* lipid synthesis for membrane formation, lipid synthesis for stress-related responses, lipid synthesis for mechanical processes in membrane trafficking etc…).

The aim of this study was therefore to investigate the possibility that some LPAATs might be associated with the functioning of the secretory pathway through PA neo-synthesis not related to the bulk neo-synthesis of membrane phospholipids.

First, we have determined that the four extra-plastidial AtLPAAT2-5 are all strict LPA acyltransferases which was an evident prerequisite for such a study. Then, we have confirmed that the AtLPAAT2,4 and 5 are ER located but determined that they do not cycle between the ER and the Golgi. AtLPAAT3 localization was unknown and, by imaging roots of plants expressing GFP-LPAAT3 and by transient expression of this construct in tobacco leaves, we have determined that AtLPAAT3 is located in ER domains and that they may correspond to ERES for some of them.

Since AtLPAAT2 is likely the primordial enzyme for the *de novo* synthesis of PA to sustain the overall *de novo* synthesis of phospholipids in the ER (Angkawijaya et al., 2017), we focused our attention to the three other AtLPAAT3-5. Given the results obtained on root growth phenotypes with the single, double and triple mutants, the strategy was then to create a combined genetic and biochemical approach as already managed successfully for the study of sphingolipids in the plant secretory pathway (Melser et al., 2010; Wattelet-Boyer et al., 2016). Treating the double mutant *lpaat4-2;lpaat5-2* with the LPAT inhibitor CI976 (Brown et al., 2008; Schmidt and Brown, 2009; Yang et al., 2011) defined experimental conditions in which a significant decrease in neo-synthesized PA without any decrease in the neo-synthesis of the major phospholipids PC and PE was obtained.

Such a result greatly suggest that under these conditions other types of LPAT (LPCAT and LPEAT which synthesize new PC and PE from lysophosphatidylcholine and lysophosphatidylethanolamine), that are present in all lines (double mutant *lpaat4-2;lpaat5-2* as well as the WT and the triple mutant *lpaat3-1;lpaat4-2;lpaat5-2*), are not greatly affected under these experimental conditions. The effect of the drug was therefore likely related to its action on the LPAATs. In addition, both AtLPAAT3 and AtLPAAT2 were the targets of the drug in the CI976_treated double mutant *lpaat4-2;lpaat5-2*, but since the major phospholipids PC and PE were not decreased, it is likely that the amount of PA synthesized was still sufficient for sustaining phospholipid synthesis but not sufficient for its role related to the transport of PIP2;7 and PIN2. Therefore, the effects observed on primary root growth and protein transport in the secretory pathway could likely be attributed to the inhibition of “specific” pool(s) of PA and that a threshold concentration of PA was required for the transport of some proteins being processed efficiently.

The fact that PIN2 was found partially retained at the TGN but not significantly at the ERES is intriguing compared to the potential localization of AtLPAAT3 in this ER domain. A first possibility could be that some of these AtLPAATs are present in the TGN but at a concentration that was not detected/detectable in our approach. In addition, AtLPAAT3 localization seemed to depend on active ER export motifs which could also support the hypothesis of its presence in a post-ER compartment. However, none of the AtLPAAT2-5 has been identified so far in Golgi/TGN proteomes (Drakakaki et al., 2012; Parsons et al., 2012; Groen et al., 2014; Heard et al., 2015), and therefore such a hypothesis can be considered unlikely for the moment with these proteomic data. Another possibility would be that ER-TGN connection may eventually support the feeding of PA to the TGN but no clear relationship has been demonstrated yet between these two compartments in plant cells as shown in mammalian cells (Mesmin et al., 2017). Anyway, these two aspects have still to be kept in mind because they may help explaining the partial retention of PIN2 at the level of the TGN.

As shown in Fig. 9, the potential impact of LPAATs on PIP2;7 transport to the plasma membrane was evidenced through both a decrease in PIP2;7 in the plasma membrane and an increase in the cytoplasm. Interestingly, PIP2;7 being highly regulated under salt stress (Pou et al., 2016) and *AtLPAAT4* gene expression being enhanced in roots under these conditions (Supplementary Fig. S9), this allowed us to perform critical experiments to strongly support our conclusion that PIP2;7 transport to the plasma membrane was linked to LPAAT activities: (i) the double mutant *lpaat4-2;lpaat5-2* was effectively more sensitive to salt stress than the WT and the triple mutant *lpaat3-1;lpaat4-2;lpaat5-2* (Fig. 10), (ii) we found that the double mutant *lpaat4-2;lpaat5-2* became less sensitive to salt stress in the presence of CI976 (Fig. 10), (iii) PIP2;7 localized less to the plasma membrane of double mutant *lpaat4-2;lpaat5-2* root cells under CI976 treatment (Fig. 9), (iv) complementation of the CI976-treated double mutant *lpaat4-2;lpaat5-2* by overexpression of *AtLPAAT4* partially restored the localization of PIP2;7 at the plasma membrane (Fig.9). Therefore, thanks to the behavior of PIP2;7 during salt stress conditions, we were able to clearly correlate the efficiency of its transport to the plasma membrane with the functionality of LPAATs. Li et al. (2019) have shown, by using a PA-specific optogenetic biosensor which determines the precise spatio-temporal dynamics of PA at the plasma membrane, that salt stress triggered an accumulation of PA via the activity of a phospholipase Dα1 (PLDα1). Investigating a pldα1 mutant indicated that PA signalling integrates with cellular pH dynamics to mediate plant response to salt stress (Li et al., 2019). Therefore, it is likely that PA is involved in both signalling and mecanistic processes in regulating various fundamental biological functions in plants.

Unfortunately, because the immunostaining strategy was not possible with PIP2;7, we could not address the question of the nature of the compartments where PIP2;7 was retained. In any case, Hachez et al. (2014) have found that PIP2;7 interacts with the SNAREs SYP61 and SYP121 to reach the plasma membrane, and we have shown that the sorting of PIN2 at the TGN occurs at a SYP61 TGN sub-domain (Wattelet-Boyer et al., 2016) where we found PIN2 partially retained. We may therefore hypothesize that PIP2;7 was to some extent retained in the same compartment as PIN2. In addition, it has been shown that under salt stress, loss of phospholipase D (PLD) function impaired auxin redistribution and this resulted in decreased primary root growth (Wang et al., 2019). Therefore, these plasma membrane proteins may have similar dependencies on some aspects of the trafficking machinery, and may also be similarly dependent on a specific formation of PA. Moreover, the data of Wang et al. (2019) indicate a role of PA in coupling extracellular salt signaling to PA-regulated PINOID kinase dependent PIN2 phosphorylation and polar auxin transport. In conclusion, PA produced by different enzymes (PLDs, LPAATs…) at different intracellular sites (early secretory pathway, late secretory pathway, plasma membrane) may regulate diverse aspects of protein trafficking, dynamics and lipid signaling functions. The recent implication of AtLPAAT4 and AtLPAAT5 in nitrogen-starvation response (Angkawijaya et al., 2019) also illustrates how the same lipid metabolizing enzymes can be engaged in multiple different cellular functions (Pokotylo et al., 2018).

Since phospholipases and therefore lysophospholipids are involved in membrane trafficking in roots and pollen (Lee et al., 2010; Kim et al., 2011), the fact that we have found an implication of AtLPAATs in the trafficking of PIP2;7 and PIN2 and primary root growth, also indicates that an interplay between phospholipases and LPAATs is likely in plant cells as already demonstrated in mammalian cells (Yang et al., 2011). Pagliuso et al. (2016) have identified a key component (CtBP1-S/BARS) of a protein complex that is required for fission of several endomembranes in mammalian cells which binds to and activates a trans-Golgi LPAAT, and that this interaction is essential for fission of transport vesicles. Interconversion of LPA and PA probably facilitates the fission process either directly or indirectly (through binding of protein(s) of the machinery to PA). In addition, the production of PA by the phospholipase D can also be critical for membrane trafficking in plant cells (Pleskot et al., 2012) as evidenced in mammalian cells for vesicle fission (Yang et al., 2011), suggesting that PA produced by different enzymes (phospholipase D, LPAATs) can be involved at different steps or pathways. We must also consider another potential complexity since some enzymes (cytosolic or membranous) may be active both as phospholipase (producing lysophospholipids from phospholipids) and acyltransferase (to reform a phospholipid) in order to contribute via lipid remodeling (Ghosh et al., 2009; Jasieniecka-Gazarkiewicz et al., 2016) to membrane deformation/re-arrangements involved in the fusion/fission processes.

In addition, since the transport of H+ATPases to the plasma membrane was not affected, this suggests either different sensitivities of the trafficking process of different proteins to PA concentration or different molecular requirements for their trafficking. Such a difference could be related to what has been observed in the role of PA in pollen tube growth (Pleskot et al., 2012) that strengthens the fact that several different mechanisms/pathways need to be considered in the complexity of the regulation of protein trafficking.

Finally, since other lipid families (sterols, sphingolipids…) are also critical in the functioning and regulation of the plant secretory pathway (Laloi et al., 2007; Men et al., 2008; Boutté et al., 2010; Melser et al., 2010; Markham et al., 2011; Wattelet-Boyer et al., 2016), we must consider that a huge interplay between lipids, lipid-synthesizing/modifying enzymes and lipid-binding proteins is at work to govern and regulate the plant secretory pathway.

As a conclusion, we have designed an experimental set up which allowed investigating the potential involvement of AtLPAATs and PA in the functioning of the plant root secretory pathway. The double mutant *lpaat4-2;lpaat5-2* treated with the LPAT inhibitor CI976 was significantly affected in primary root growth and the trafficking of PIP2;7 and PIN2 was found to be disturbed. Our results support a critical PA concentration threshold involved in the transport of some proteins through the plant root secretory pathway. Since phospholipases and therefore lysophospholipids are involved in protein membrane trafficking in roots (Lee et al., 2010), the implication of AtLPAATs in the trafficking of PIP2;7 and PIN2 in roots also suggests an interplay between phospholipases and LPAATs in root cells as shown in mammalian cells (Yang et al., 2011).

## Supplementary data

**Supplementary Table 1**. Primers used in the study

**Supplementary Fig. S1** Alignment of the Human LPAAT3 with the AtLPAATs 2-5 from

*Arabidopsis*

**Supplementary Fig. S2** *AtLPAAT3, AtLPAAT4* and *AtLPAAT5* are expressed at similar levels in *Arabidospis* roots

**Supplementary Fig. S3** Caracterization of *lpaat* insertion mutant lines and T-DNA positions

**Supplementary Fig. S4** Primary root length of the triple mutant *lpaat3-1;lpaat4-2;lpaat5-2* is stimulated

**Supplementary Fig. S5** *AtLPAAT2* expression level in primary roots

**Supplementary Fig. S6** Trafficking of PM H+ ATPases is not altered upon CI976 treatment

**Supplementary Fig. S7** PIN2-GFP accumulates in punctuated structures that colocalise with a TGN marker in CI976-treated *lpaat4-2,lpaat5-2* mutant

**Supplementary Fig. S8** Salt stress (NaCl 150 mM) impact on *LPAAT4* gene expression in roots

**Supplementary Fig. S9** Expression of *AtLPAAT4* in 5-days old *Arabidopsis* roots

**Supplementary Fig. S10**. Endocytosis is not accelerated in the double mutant *lpaat4-2;lpaat5-2* line treated with 10µM of CI-976

**Supplementary Fig. S11** Semi-quantitative RT-PCR analysis of At*LPAAT*s transcripts in roots from 5-days old plants

## Acknowledgements

This work was supported by the CNRS (Centre National de la Recherche Scientifique) and the University of Bordeaux. We thank the Bordeaux Imaging Center, part of the National Infrastructure France-BioImaging supported by the French National Research Agency (ANR-10-INBS-04).

## Author contributions

V.W-B., M.L.G., Y.B., J-J.B and P.M. conceived the different parts of the study; V.W-B. performed most of the experiments; M.L.G. performed the *in vitro* LPAAT activities in *E*.*coli*; F. D-D. and L. M-P. performed the assays on the effect of CI976 on phospholipid *de-novo* synthesis; V.K. performed the colocalization of LPAAT3 with SAR1. V.W-B., Y.B. and P.M. analyzed the microscopy data; V.W-B., J-J.B and P.M. analyzed all the data and P.M. supervised the experiments; P.M. wrote the article with contributions of all the authors; P.M. is the author responsible for contact and ensures communication.

## Data availability statement

The data supporting the findings of this study are available within the paper and within its supplementary data.

## Notes

### Competing Interest Statement

The authors have declared no competing interest.

