## Supplementary Figures 1-11 and supplementary Table 1 for "Lyso-Phosphatidic Acid Acyl-Transferases: a link with intracellular protein transport in *Arabidopsis* root cells?": Supplementary Data Wattelet-Boyer et al.pptx

### Slide 1
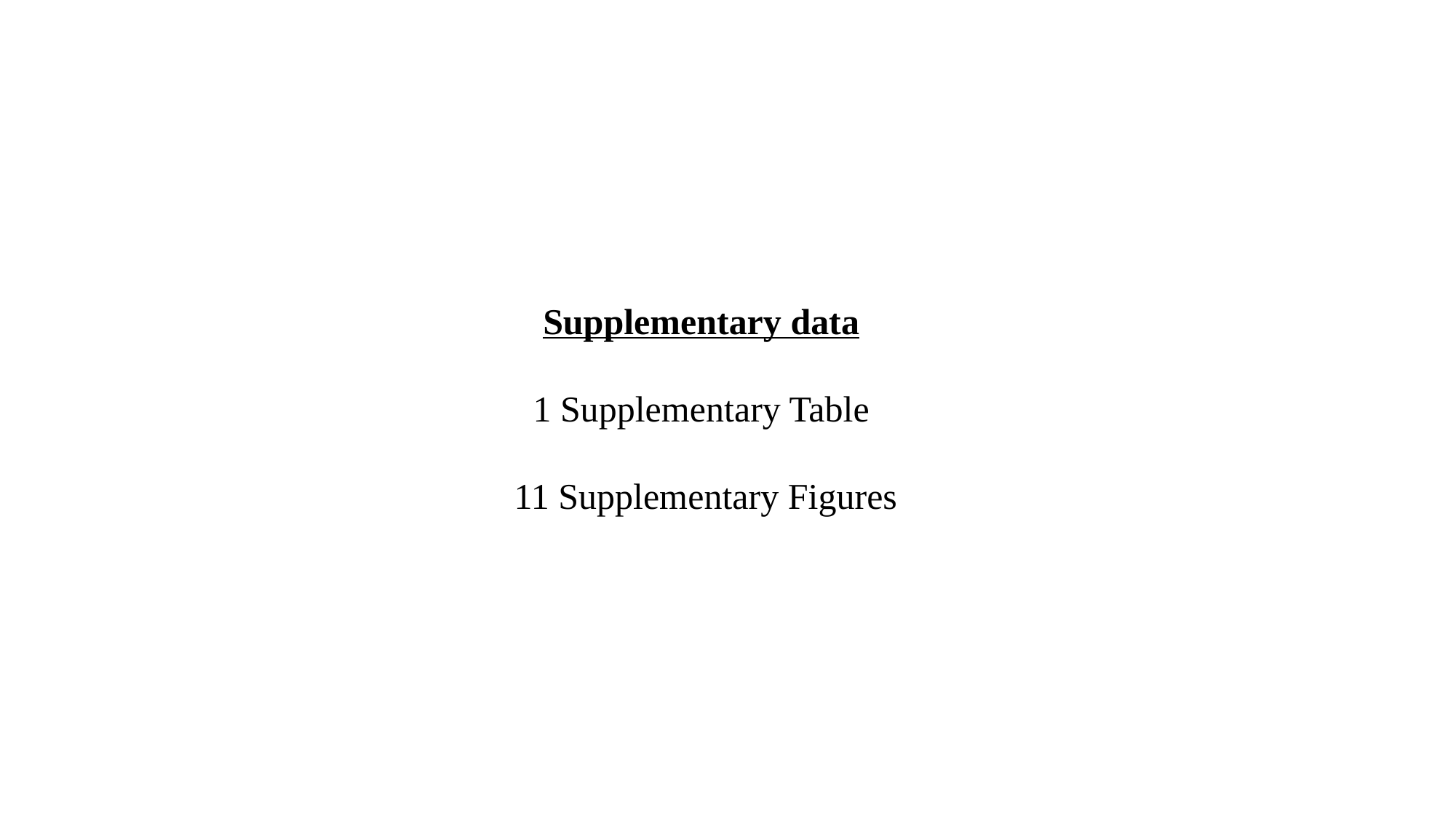

Supplementary data
1 Supplementary Table
11 Supplementary Figures

### Slide 2
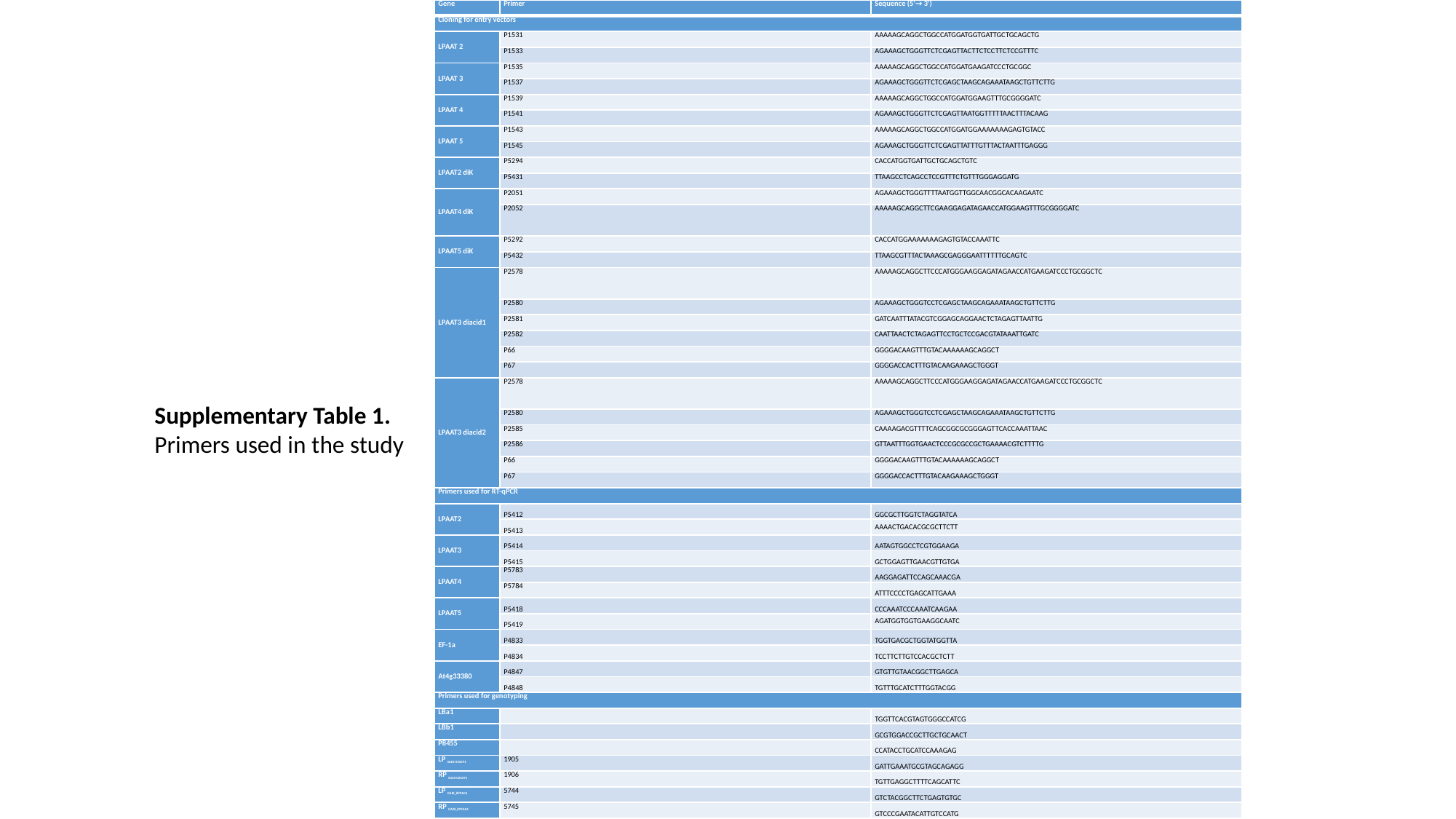

| Gene | Primer | Sequence (5’→ 3’) |
| --- | --- | --- |
| Cloning for entry vectors | | |
| LPAAT 2 | P1531 | AAAAAGCAGGCTGGCCATGGATGGTGATTGCTGCAGCTG |
| | P1533 | AGAAAGCTGGGTTCTCGAGTTACTTCTCCTTCTCCGTTTC |
| LPAAT 3 | P1535 | AAAAAGCAGGCTGGCCATGGATGAAGATCCCTGCGGC |
| | P1537 | AGAAAGCTGGGTTCTCGAGCTAAGCAGAAATAAGCTGTTCTTG |
| LPAAT 4 | P1539 | AAAAAGCAGGCTGGCCATGGATGGAAGTTTGCGGGGATC |
| | P1541 | AGAAAGCTGGGTTCTCGAGTTAATGGTTTTTAACTTTACAAG |
| LPAAT 5 | P1543 | AAAAAGCAGGCTGGCCATGGATGGAAAAAAAGAGTGTACC |
| | P1545 | AGAAAGCTGGGTTCTCGAGTTATTTGTTTACTAATTTGAGGG |
| LPAAT2 diK | P5294 | CACCATGGTGATTGCTGCAGCTGTC |
| | P5431 | TTAAGCCTCAGCCTCCGTTTCTGTTTGGGAGGATG |
| LPAAT4 diK | P2051 | AGAAAGCTGGGTTTTAATGGTTGGCAACGGCACAAGAATC |
| | P2052 | AAAAAGCAGGCTTCGAAGGAGATAGAACCATGGAAGTTTGCGGGGATC |
| LPAAT5 diK | P5292 | CACCATGGAAAAAAAGAGTGTACCAAATTC |
| | P5432 | TTAAGCGTTTACTAAAGCGAGGGAATTTTTTGCAGTC |
| LPAAT3 diacid1 | P2578 | AAAAAGCAGGCTTCCCATGGGAAGGAGATAGAACCATGAAGATCCCTGCGGCTC |
| | P2580 | AGAAAGCTGGGTCCTCGAGCTAAGCAGAAATAAGCTGTTCTTG |
| | P2581 | GATCAATTTATACGTCGGAGCAGGAACTCTAGAGTTAATTG |
| | P2582 | CAATTAACTCTAGAGTTCCTGCTCCGACGTATAAATTGATC |
| | P66 | GGGGACAAGTTTGTACAAAAAAGCAGGCT |
| | P67 | GGGGACCACTTTGTACAAGAAAGCTGGGT |
| LPAAT3 diacid2 | P2578 | AAAAAGCAGGCTTCCCATGGGAAGGAGATAGAACCATGAAGATCCCTGCGGCTC |
| | P2580 | AGAAAGCTGGGTCCTCGAGCTAAGCAGAAATAAGCTGTTCTTG |
| | P2585 | CAAAAGACGTTTTCAGCGGCGCGGGAGTTCACCAAATTAAC |
| | P2586 | GTTAATTTGGTGAACTCCCGCGCCGCTGAAAACGTCTTTTG |
| | P66 | GGGGACAAGTTTGTACAAAAAAGCAGGCT |
| | P67 | GGGGACCACTTTGTACAAGAAAGCTGGGT |
| Primers used for RT-qPCR | | |
| LPAAT2 | P5412 | GGCGCTTGGTCTAGGTATCA |
| | P5413 | AAAACTGACACGCGCTTCTT |
| LPAAT3 | P5414 | AATAGTGGCCTCGTGGAAGA |
| | P5415 | GCTGGAGTTGAACGTTGTGA |
| LPAAT4 | P5783 | AAGGAGATTCCAGCAAACGA |
| | P5784 | ATTTCCCCTGAGCATTGAAA |
| LPAAT5 | P5418 | CCCAAATCCCAAATCAAGAA |
| | P5419 | AGATGGTGGTGAAGGCAATC |
| EF-1a | P4833 | TGGTGACGCTGGTATGGTTA |
| | P4834 | TCCTTCTTGTCCACGCTCTT |
| At4g33380 | P4847 | GTGTTGTAACGGCTTGAGCA |
| | P4848 | TGTTTGCATCTTTGGTACGG |
| Primers used for genotyping | | |
| LBa1 | | TGGTTCACGTAGTGGGCCATCG |
| LBb1 | | GCGTGGACCGCTTGCTGCAACT |
| P8455 | | CCATACCTGCATCCAAAGAG |
| LP _SALK-020291_ | 1905 | GATTGAAATGCGTAGCAGAGG |
| RP _SALK-020291_ | 1906 | TGTTGAGGCTTTTCAGCATTC |
| LP _GABI\_899A04_ | 5744 | GTCTACGGCTTCTGAGTGTGC |
| RP _GABI\_899A04_ | 5745 | GTCCCGAATACATTGTCCATG |
Supplementary Table 1.
Primers used in the study

### Slide 3
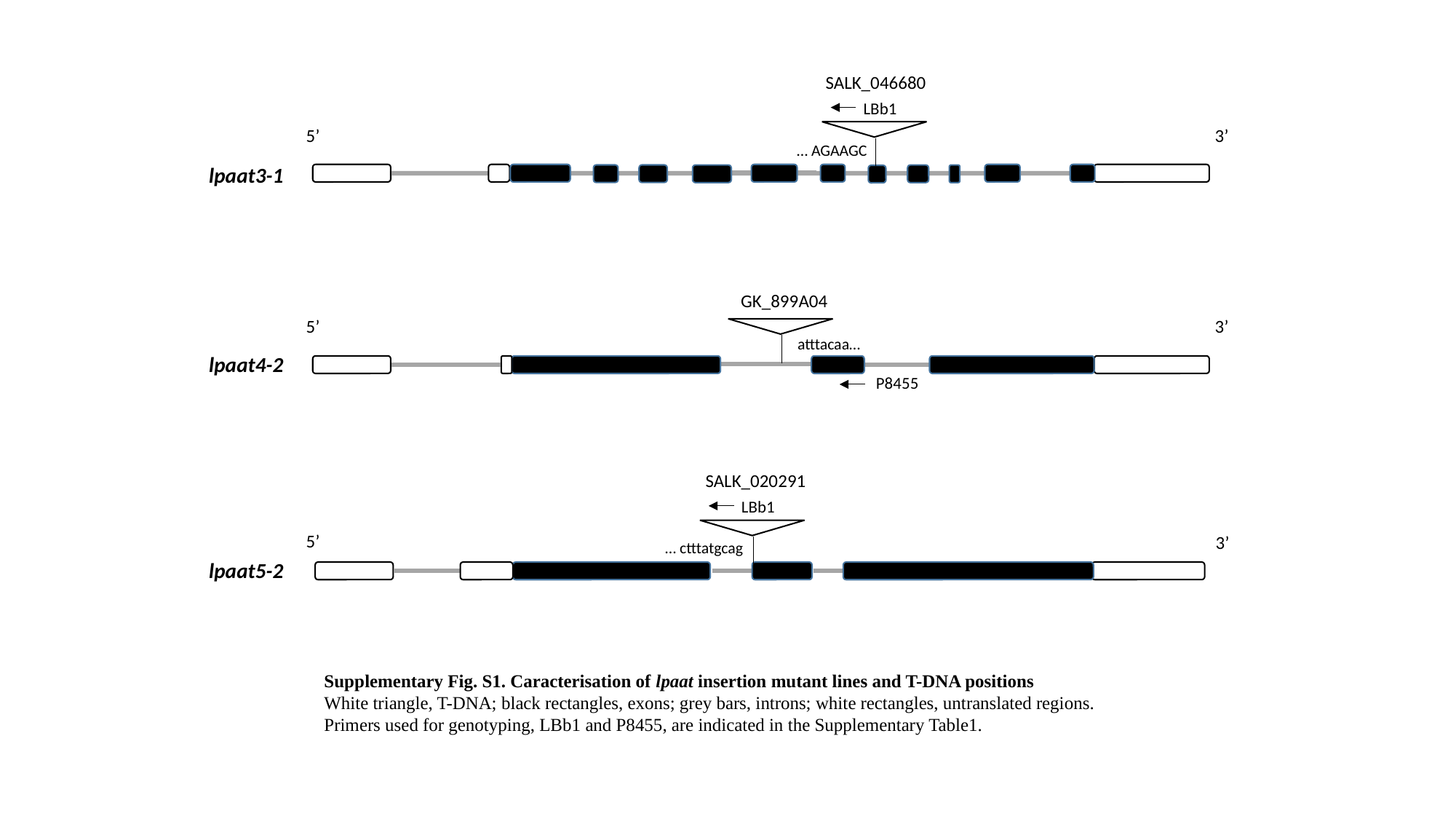

SALK_046680
LBb1
5’
3’
… AGAAGC
lpaat3-1
GK_899A04
5’
3’
atttacaa…
lpaat4-2
P8455
SALK_020291
LBb1
5’
3’
… ctttatgcag
lpaat5-2
Supplementary Fig. S1. Caracterisation of lpaat insertion mutant lines and T-DNA positions
White triangle, T-DNA; black rectangles, exons; grey bars, introns; white rectangles, untranslated regions.
Primers used for genotyping, LBb1 and P8455, are indicated in the Supplementary Table1.

### Slide 4
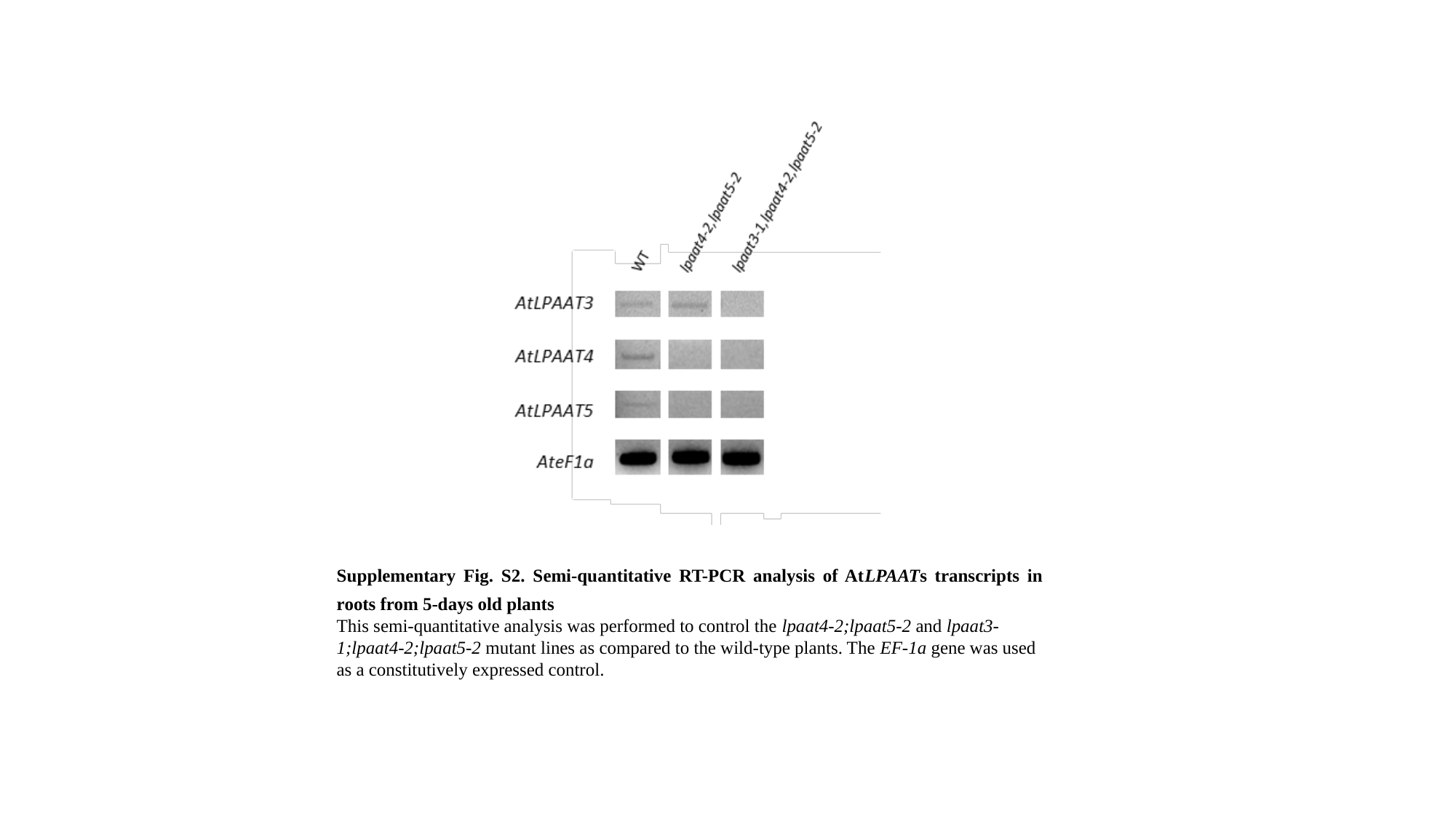

Supplementary Fig. S2. Semi-quantitative RT-PCR analysis of AtLPAATs transcripts in roots from 5-days old plants
This semi-quantitative analysis was performed to control the lpaat4-2;lpaat5-2 and lpaat3-1;lpaat4-2;lpaat5-2 mutant lines as compared to the wild-type plants. The EF-1a gene was used as a constitutively expressed control.

### Slide 5
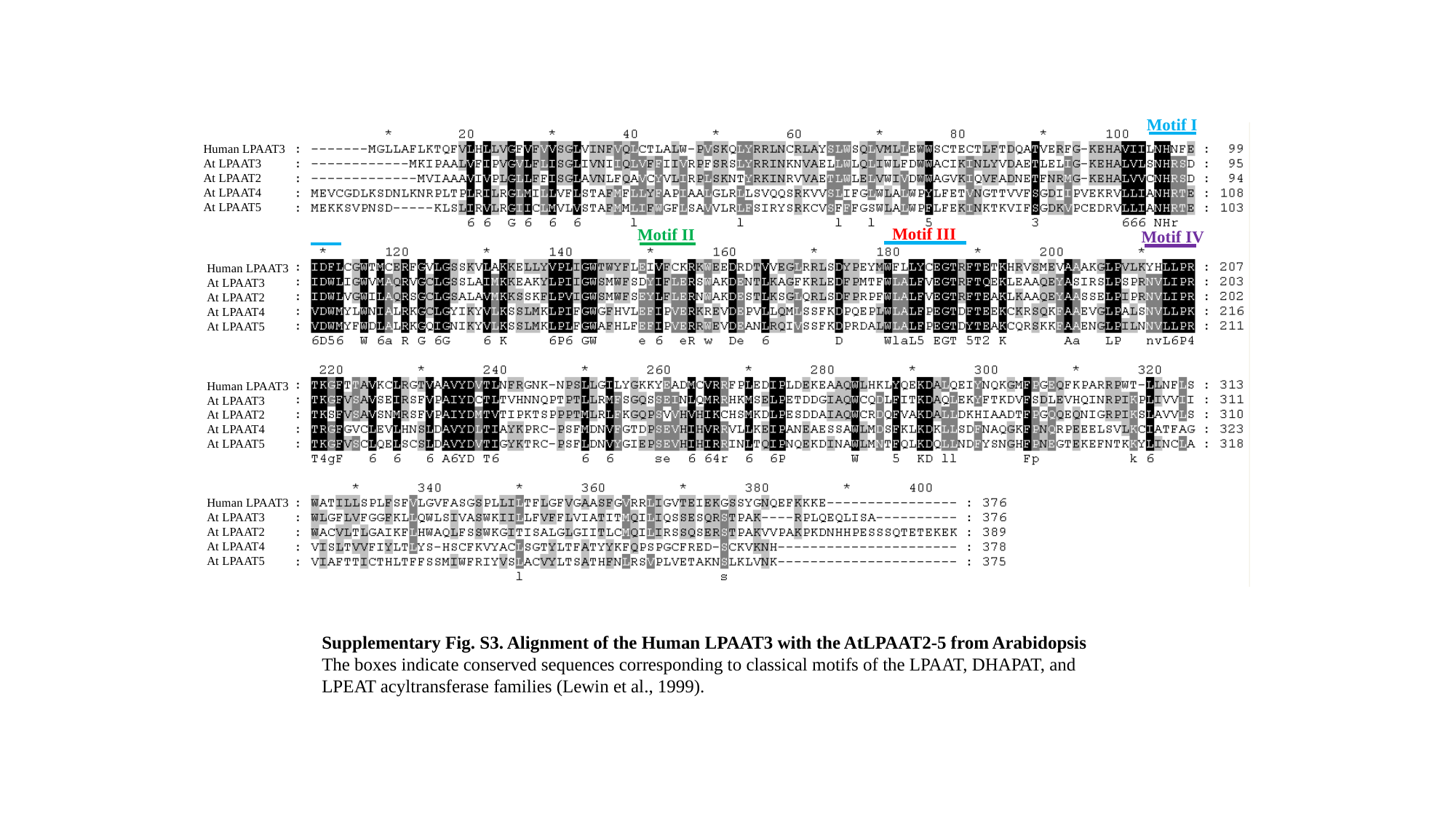

Motif I
Human LPAAT3
At LPAAT3
At LPAAT2
At LPAAT4
At LPAAT5
Motif III
Motif II
Motif IV
Human LPAAT3
At LPAAT3
At LPAAT2
At LPAAT4
At LPAAT5
Human LPAAT3
At LPAAT3
At LPAAT2
At LPAAT4
At LPAAT5
Human LPAAT3
At LPAAT3
At LPAAT2
At LPAAT4
At LPAAT5
Supplementary Fig. S3. Alignment of the Human LPAAT3 with the AtLPAAT2-5 from Arabidopsis
The boxes indicate conserved sequences corresponding to classical motifs of the LPAAT, DHAPAT, and
LPEAT acyltransferase families (Lewin et al., 1999).

### Slide 6
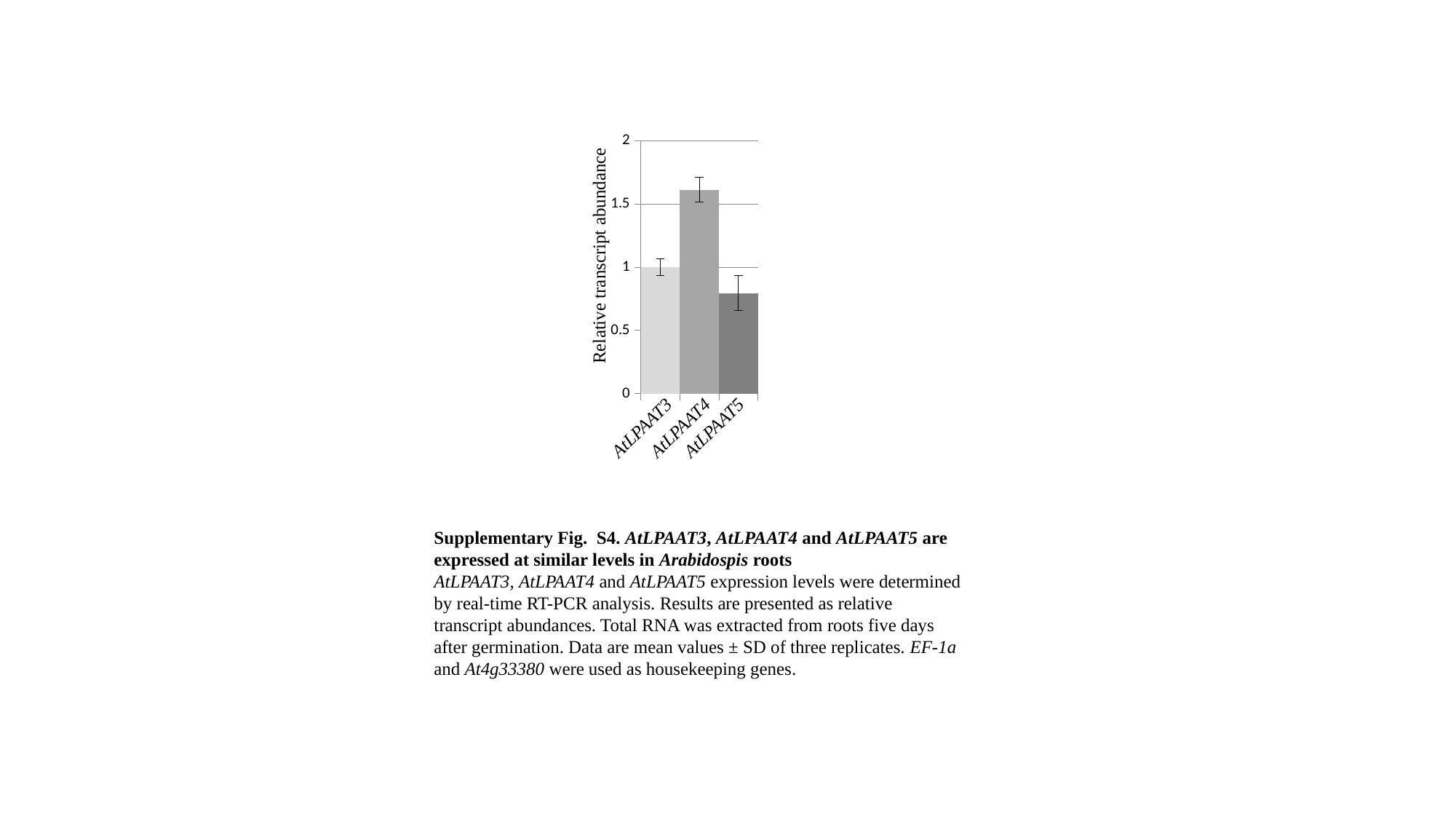

#### Chart
| Category | |
|---|---|
| LPAAT3 | 1.0 |
| LPAAT4 | 1.6132835184442542 |
| LPAAT5 | 0.7955364837549151 |Relative transcript abundance
AtLPAAT3
AtLPAAT4
AtLPAAT5
Supplementary Fig. S4. AtLPAAT3, AtLPAAT4 and AtLPAAT5 are expressed at similar levels in Arabidospis roots
AtLPAAT3, AtLPAAT4 and AtLPAAT5 expression levels were determined by real-time RT-PCR analysis. Results are presented as relative transcript abundances. Total RNA was extracted from roots five days after germination. Data are mean values ± SD of three replicates. EF-1a and At4g33380 were used as housekeeping genes.

### Slide 7
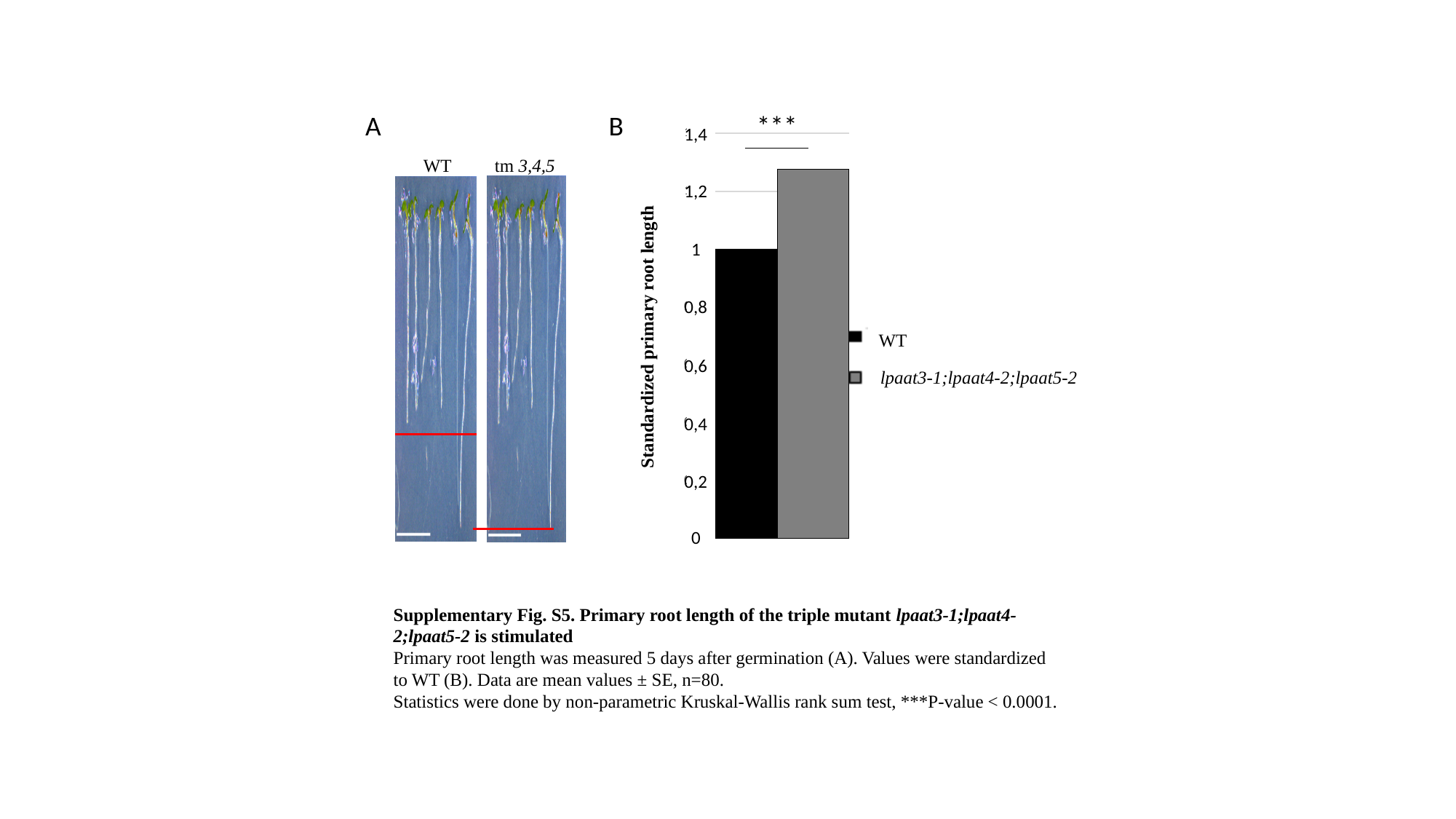

#### Chart
| Category |
|---|
WT
lpaat3-1;lpaat4-2;lpaat5-2
Standardized primary root length
1,4
1,2
1
0,8
0,6
0,4
0,2
0
#### Chart
| Category | |
|---|---|
| WT | 1.0 |
| tm | 1.276262111399585 |
***
A
B
WT
tm 3,4,5
Supplementary Fig. S5. Primary root length of the triple mutant lpaat3-1;lpaat4-2;lpaat5-2 is stimulated
Primary root length was measured 5 days after germination (A). Values were standardized to WT (B). Data are mean values ± SE, n=80.
Statistics were done by non-parametric Kruskal-Wallis rank sum test, ***P-value < 0.0001.

### Slide 8
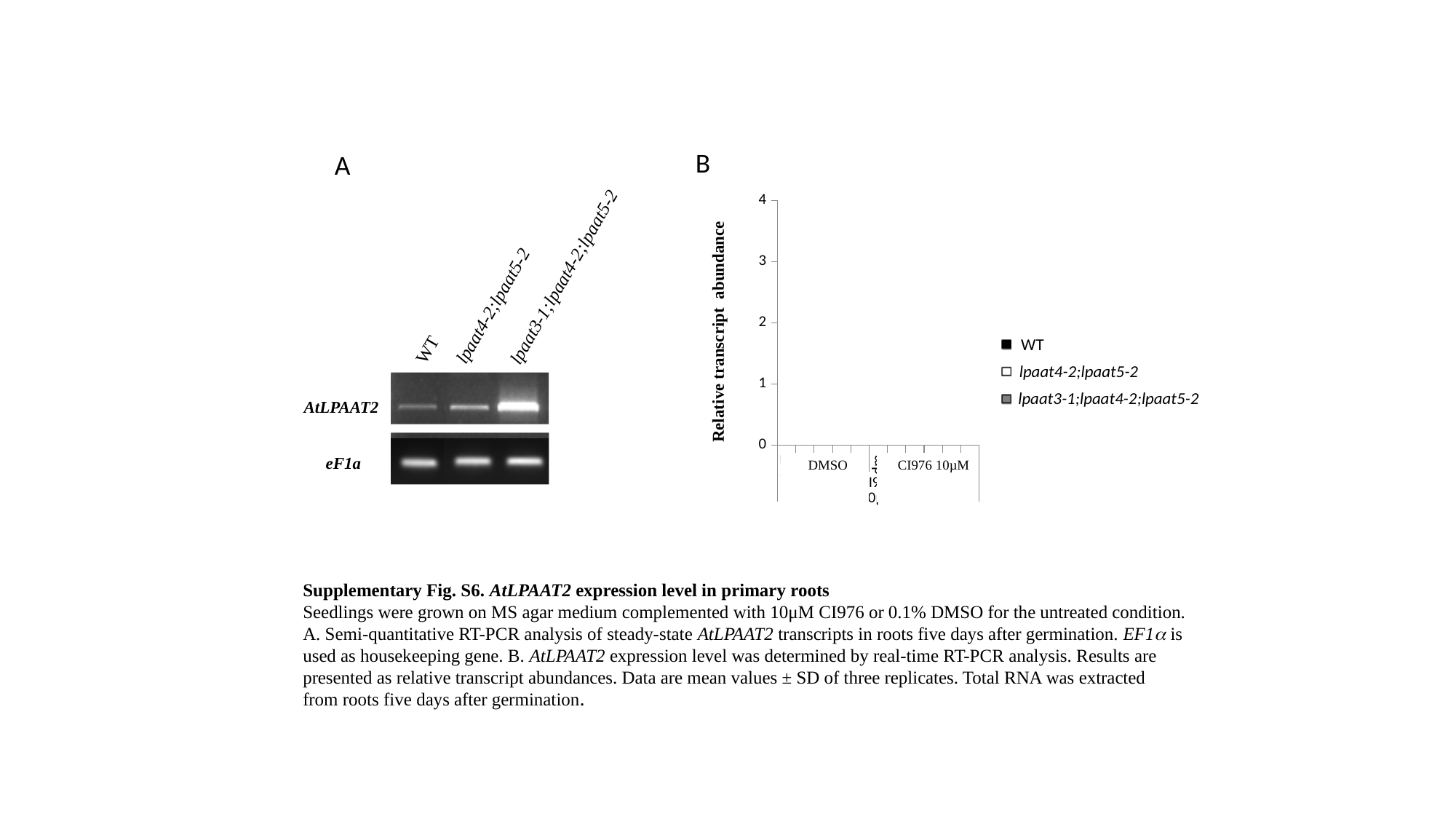

B
#### Chart
| Category | |
|---|---|
| WT | 1.0 |
| dm | 0.9383277065256009 |
| tm | 2.8951475661812873 |
| | 0.0 |
| WT | 2.0506493226237232 |
| dm | 2.1761455921185027 |
| tm | 3.3153427384239627 |
Relative transcript abundance
A
lpaat3-1;lpaat4-2;lpaat5-2
lpaat4-2;lpaat5-2
WT
AtLPAAT2
eF1a
WT
lpaat4-2;lpaat5-2
lpaat3-1;lpaat4-2;lpaat5-2
DMSO
CI976 10µM
Supplementary Fig. S6. AtLPAAT2 expression level in primary roots
Seedlings were grown on MS agar medium complemented with 10μM CI976 or 0.1% DMSO for the untreated condition.
A. Semi-quantitative RT-PCR analysis of steady-state AtLPAAT2 transcripts in roots five days after germination. EF1α is
used as housekeeping gene. B. AtLPAAT2 expression level was determined by real-time RT-PCR analysis. Results are
presented as relative transcript abundances. Data are mean values ± SD of three replicates. Total RNA was extracted
from roots five days after germination.

### Slide 9
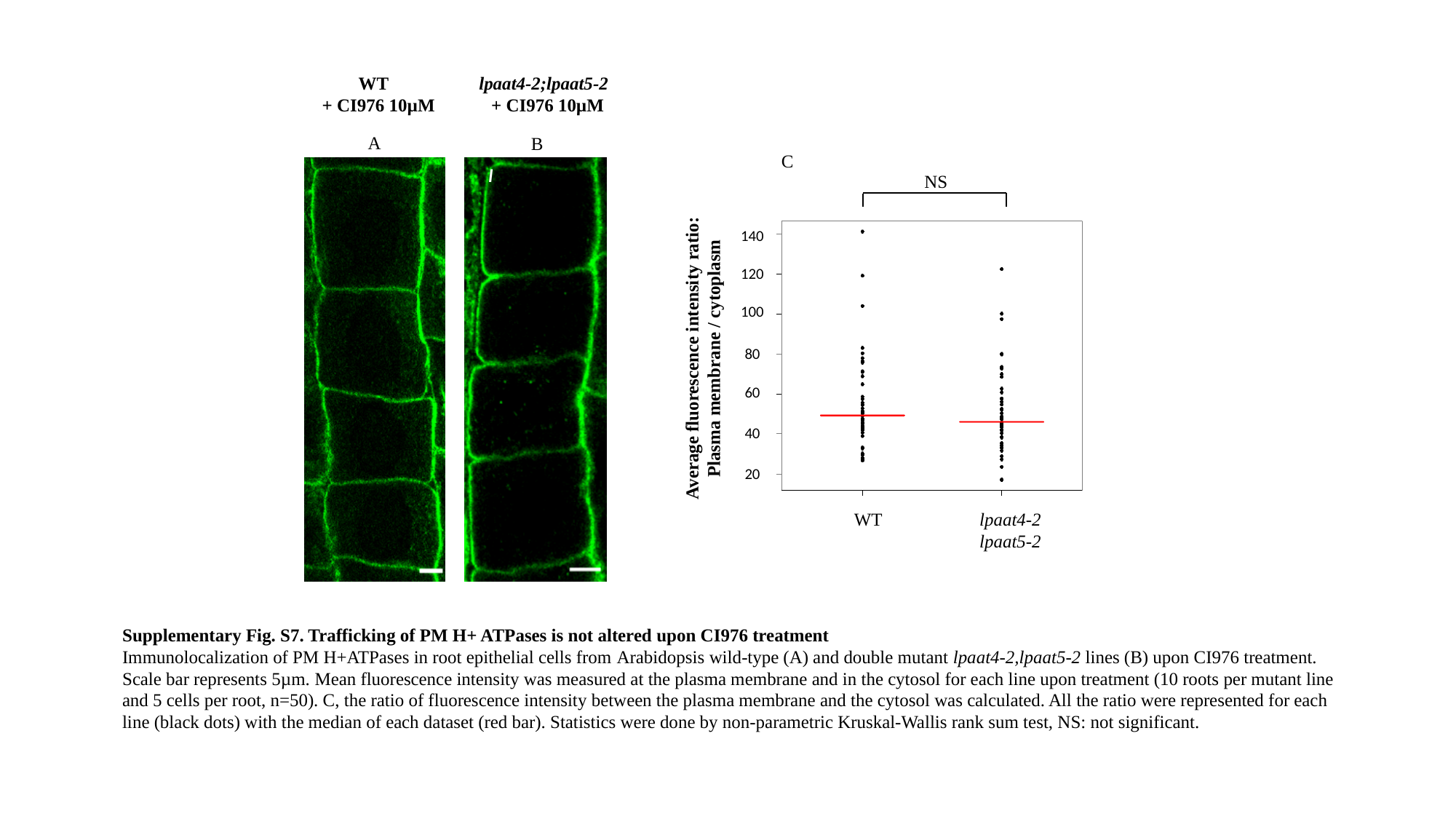

WT
+ CI976 10µM
lpaat4-2;lpaat5-2
+ CI976 10µM
A
B
C
NS
140
120
100
Average fluorescence intensity ratio:
Plasma membrane / cytoplasm
80
60
40
20
WT
lpaat4-2
lpaat5-2
l
Supplementary Fig. S7. Trafficking of PM H+ ATPases is not altered upon CI976 treatment
Immunolocalization of PM H+ATPases in root epithelial cells from Arabidopsis wild-type (A) and double mutant lpaat4-2,lpaat5-2 lines (B) upon CI976 treatment. Scale bar represents 5µm. Mean fluorescence intensity was measured at the plasma membrane and in the cytosol for each line upon treatment (10 roots per mutant line and 5 cells per root, n=50). C, the ratio of fluorescence intensity between the plasma membrane and the cytosol was calculated. All the ratio were represented for each line (black dots) with the median of each dataset (red bar). Statistics were done by non-parametric Kruskal-Wallis rank sum test, NS: not significant.

### Slide 10
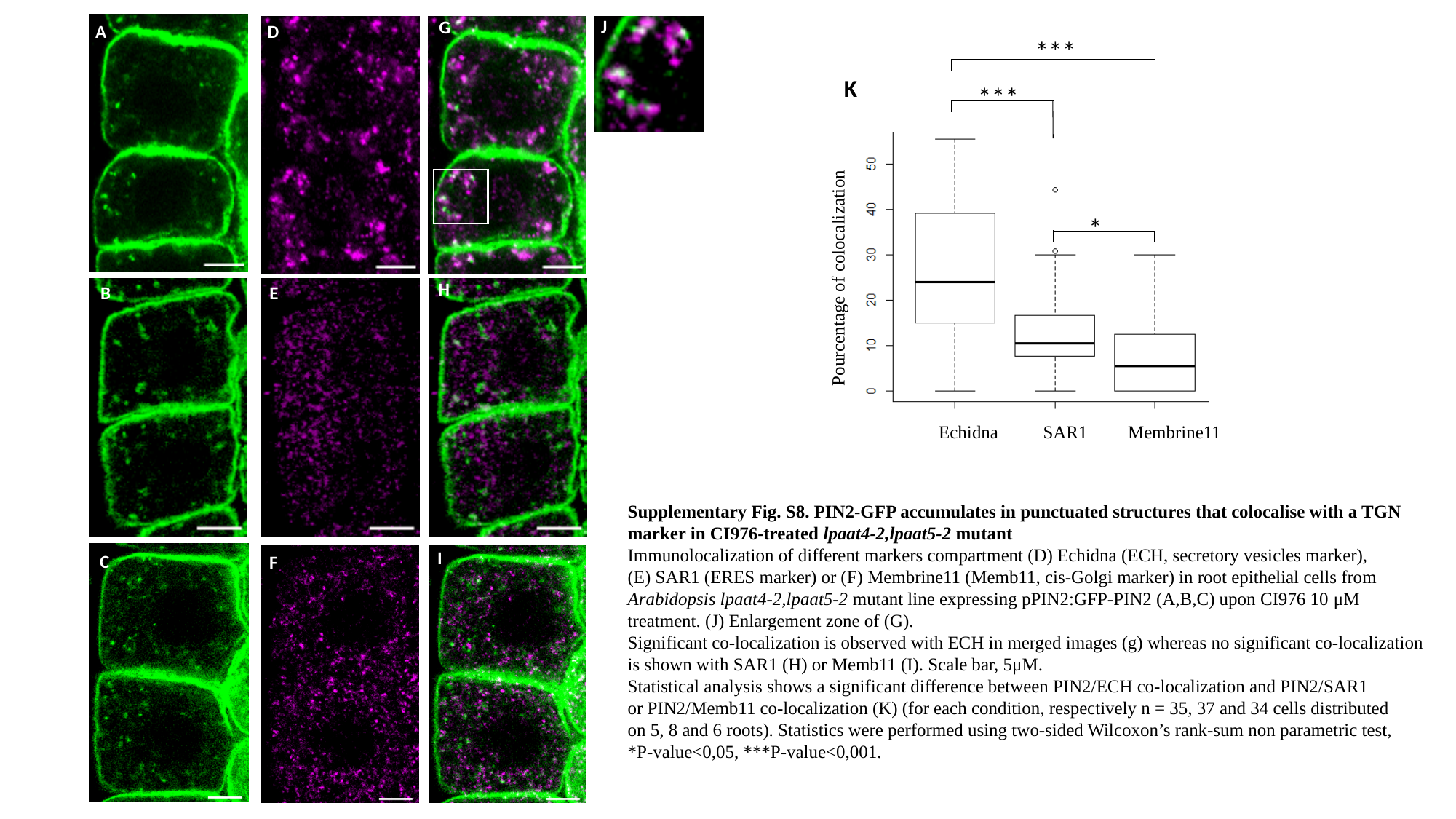

J
j
G
A
D
B
E
C
F
H
I
***
***
*
Pourcentage of colocalization
K
Echidna
SAR1
Membrine11
Supplementary Fig. S8. PIN2-GFP accumulates in punctuated structures that colocalise with a TGN marker in CI976-treated lpaat4-2,lpaat5-2 mutant
Immunolocalization of different markers compartment (D) Echidna (ECH, secretory vesicles marker),
(E) SAR1 (ERES marker) or (F) Membrine11 (Memb11, cis-Golgi marker) in root epithelial cells from
Arabidopsis lpaat4-2,lpaat5-2 mutant line expressing pPIN2:GFP-PIN2 (A,B,C) upon CI976 10 μM treatment. (J) Enlargement zone of (G).
Significant co-localization is observed with ECH in merged images (g) whereas no significant co-localization is shown with SAR1 (H) or Memb11 (I). Scale bar, 5μM.
Statistical analysis shows a significant difference between PIN2/ECH co-localization and PIN2/SAR1
or PIN2/Memb11 co-localization (K) (for each condition, respectively n = 35, 37 and 34 cells distributed
on 5, 8 and 6 roots). Statistics were performed using two-sided Wilcoxon’s rank-sum non parametric test,
*P-value<0,05, ***P-value<0,001.

### Slide 11
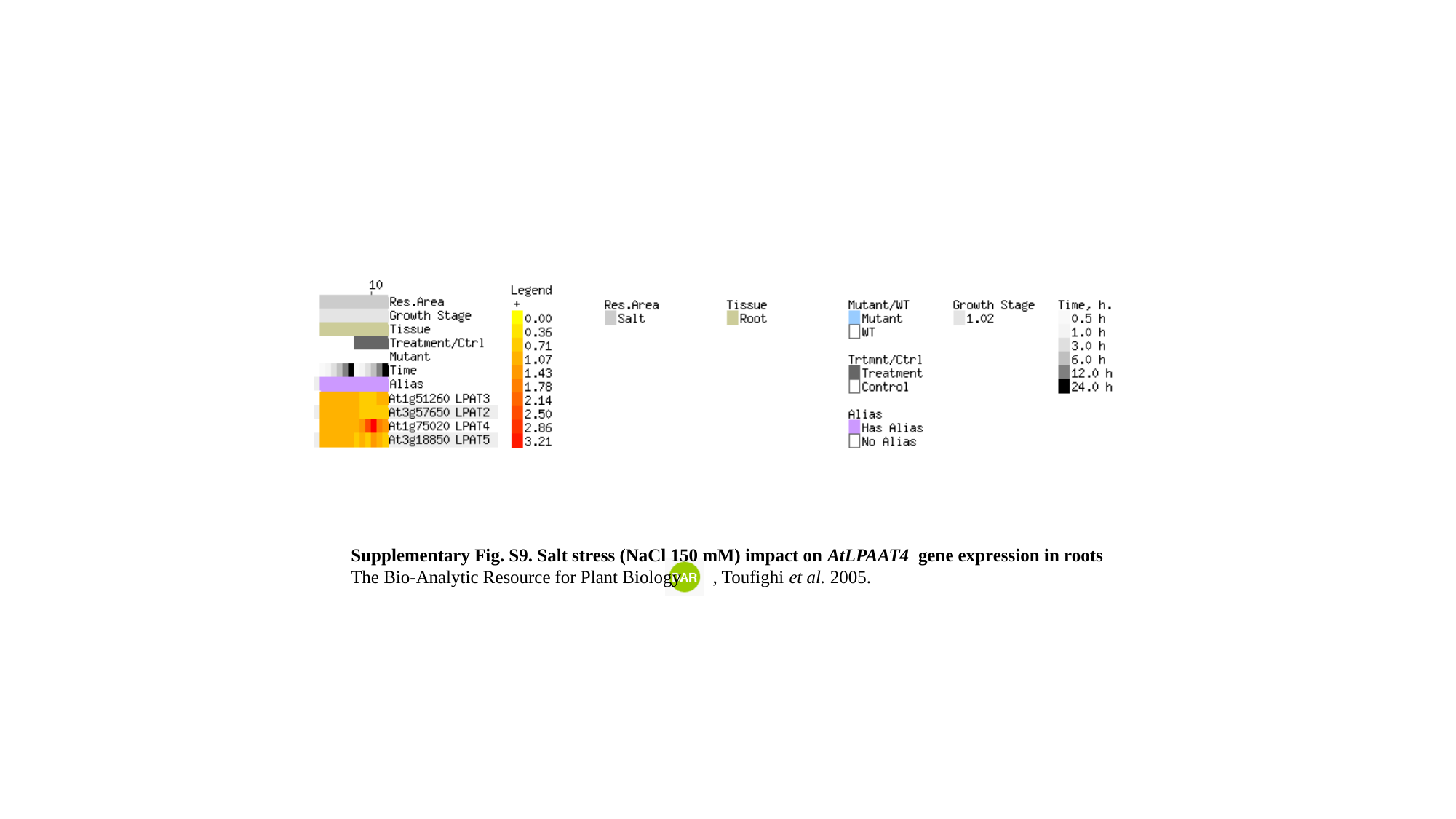

Supplementary Fig. S9. Salt stress (NaCl 150 mM) impact on AtLPAAT4 gene expression in roots
The Bio-Analytic Resource for Plant Biology , Toufighi et al. 2005.

### Slide 12
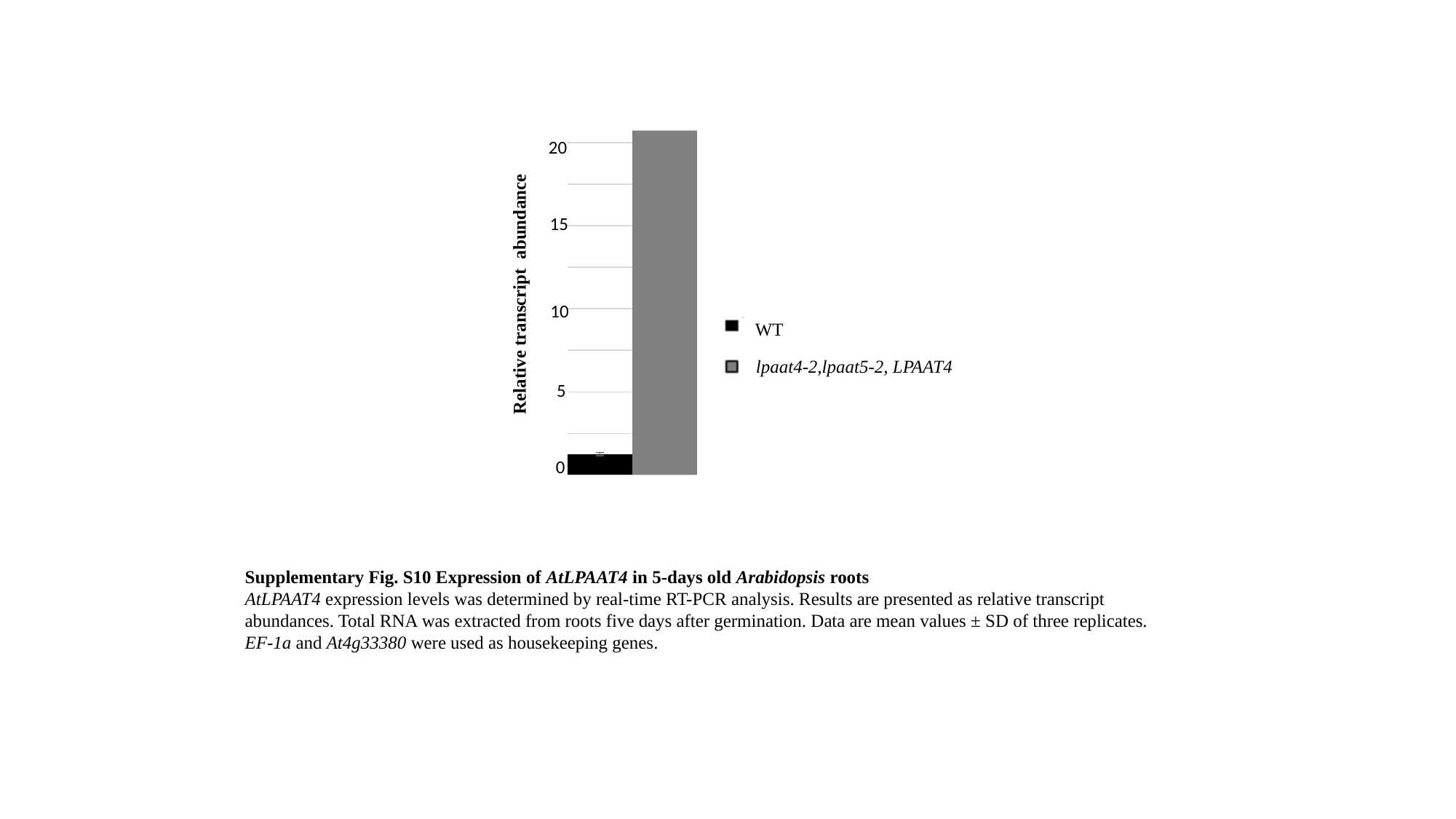

25
#### Chart
| Category | |
|---|---|
| WT | 1.0 |
| lpaat4,lpaat5 - LPAAT4 OE | 18.37613244595167 |20
15
10
5
0
Relative transcript abundance
WT
lpaat4-2,lpaat5-2, LPAAT4
Supplementary Fig. S10 Expression of AtLPAAT4 in 5-days old Arabidopsis roots
AtLPAAT4 expression levels was determined by real-time RT-PCR analysis. Results are presented as relative transcript abundances. Total RNA was extracted from roots five days after germination. Data are mean values ± SD of three replicates. EF-1a and At4g33380 were used as housekeeping genes.

### Slide 13
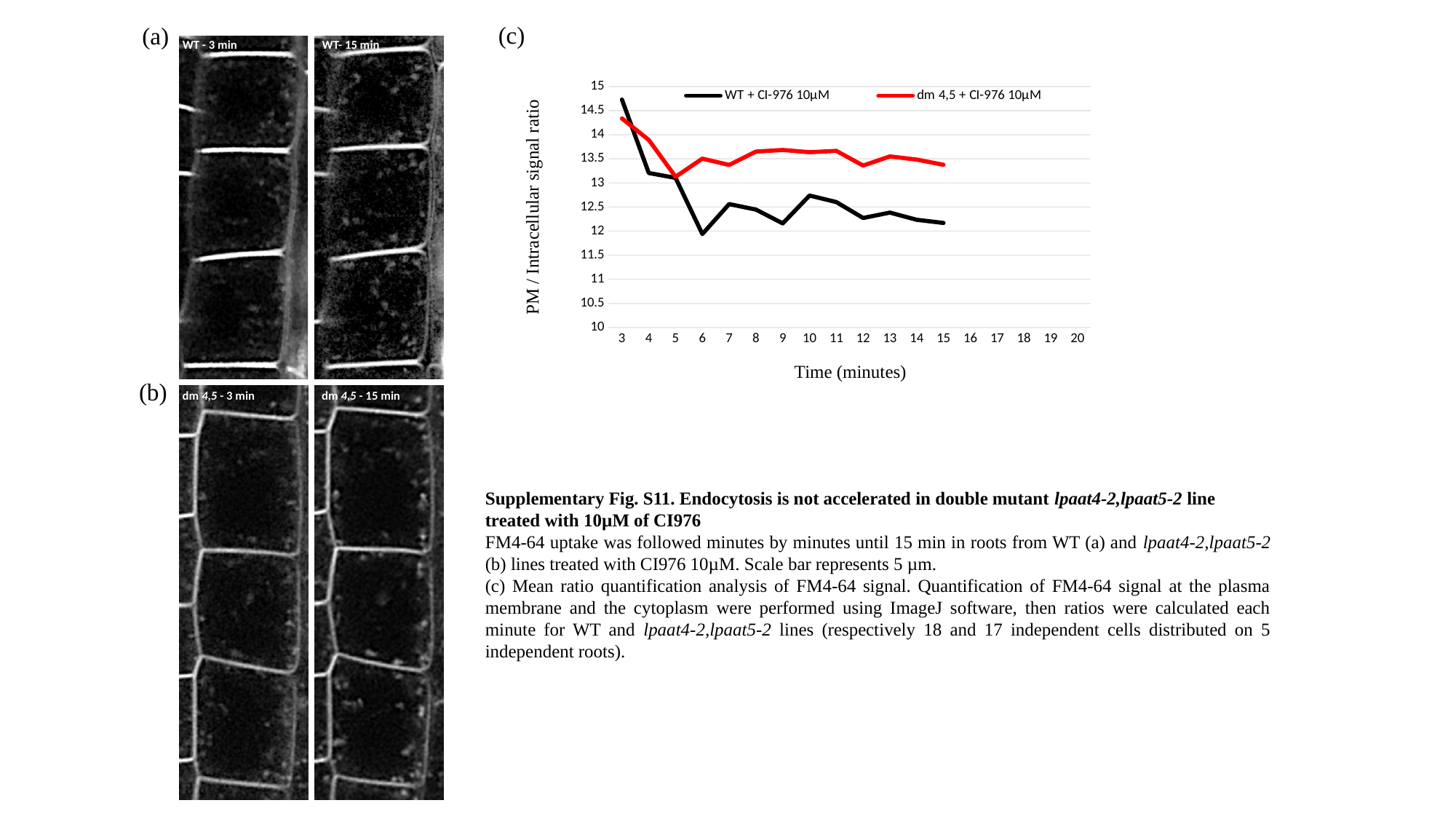

(c)
(a)
dm 4,5 - 3 min
dm 4,5 - 15 min
Time (minutes)
(b)
[unsupported chart]
PM / Intracellular signal ratio
WT - 3 min
WT- 15 min
dm 4,5 - 3 min
dm 4,5 - 15 min
WT- 15 min
Supplementary Fig. S11. Endocytosis is not accelerated in double mutant lpaat4-2,lpaat5-2 line treated with 10µM of CI976
FM4-64 uptake was followed minutes by minutes until 15 min in roots from WT (a) and lpaat4-2,lpaat5-2 (b) lines treated with CI976 10µM. Scale bar represents 5 µm.
(c) Mean ratio quantification analysis of FM4-64 signal. Quantification of FM4-64 signal at the plasma membrane and the cytoplasm were performed using ImageJ software, then ratios were calculated each minute for WT and lpaat4-2,lpaat5-2 lines (respectively 18 and 17 independent cells distributed on 5 independent roots).
